# Evaluation of metabolic and functional properties of cholesterol-reducing and GABA-producer *Limosilactobacillus fermentum* strain AGA52 isolated from lactic acid fermented Shalgam by using *in vitro* and *in silico* probiogenomic approaches

**DOI:** 10.1101/2022.10.25.513655

**Authors:** Ahmet Yetiman, Mehmet Horzum, Mikail Akbulut

## Abstract

The objective of this study is characterization of the genomic and metabolic properties of a novel *Lb. fermentum* strain AGA52 which was isolated from a lactic acid fermented beverage called “Shalgam”. The genome size of AGA52 was 2,001,084 bp, which is predicted to carry 2,024 genes, including 50 tRNAs, 3 rRNAs, 3ncRNAs, 15 CRISPR repeats, 14 CRISPR spacers, and 1 CRISPR array. The genome has a GC content of 51.82% including 95 predicted pseudogenes, 56 complete or partial transposases and 2 intact prophages. The similarity of the clusters of orthologous groups (COG) was analysed by comparison with the other *Lb. fermentum* strains. The detected resistome on the genome of AGA52 was found to be intrinsically originated. Based on genome analyses many CAZYmes were identified that are responsible for carbohydrate degradation and adaptation to gastrointestinal conditions. Besides, it has been determined that AGA52 has an obligate heterofermentative carbohydrate metabolism due to the absence of the 1-phosphofructokinase (*pfK*) enzyme. Additionally, it is determined that the strain has a higher antioxidant capacity and is adaptable to gastrointestinal simulated environments. It was also observed that the AGA52 has antimicrobial activity against *Y. Enterocolitica* ATCC9610, *B. cereus* ATCC33019, *S. enterica* sv. Typhimurium, *E*.*coli* O157:h7 ATCC43897, *L*.*monocytogenes* ATCC7644, *K*.*pneumoniae* ATCC13883, and *P. vulgaris* ATCC8427. Cholesterol assimilation (33.9±0.005%) and GABA production capacities were also confirmed by “*in silico*” and “*in vitro*”. Overall, investigation of genomic and metabolic features of the AGA52 revealed that is a potential psychobiotic and probiotic dietary supplement candidate and can bring functional benefits to the host.

## Introduction

The lactobacilli are a bacterial genus that is a member of Lactic acid bacteria (LAB) which can be found in fermented foods and the gastrointestinal microbiome of humans and animals [1]. They are widely used in the food biotech industry because of their well-documented beneficial and functional effects on health [2]. The members of the lactobacilli can produce different kinds of metabolic products such as bacteriocins and various organic acids, which generate an unsuitable environment for the growth of pathogenic and saprophytic microorganisms [3, 4]. Furthermore, metabolites or bacterial cells by themselves can show antiviral activity [5, 6]. Many species from the *Lactobacillus*, have been recognized as GRAS (Generally Recognized As Safe) and/or QPS (Presumably Qualified as Safe), due to their potential probiotic properties. Probiotics are defined as nonpathogenic living microorganisms that confer health benefits to the host when administered in adequate amounts by the International Scientific Association for Probiotics and Prebiotics (ISAPP) [7].

Lately, the genus *Lactobacillus* has been reclassified into 25 different genera, 23 of which are novel. In accordance with this new taxonomic order, *Lactobacillus fermentum* has been renamed *Limosilactobacillus fermentum* and qualified as gram-positive, coccoid, or rod-shaped, aerotolerant or anaerobic, and heterofermentative. *L. fermentum* has been generally isolated from fermented plant materials including cereals, dairy products, feces, manure, sewage, oral cavity, human vagina, and breast milk [8, 9]. Moreover, it has been reported that *L fermentum* exhibits health-promoting effects on the host when administered time-dependently. These effects are an increment of the amount of short-chain fatty acid in the colon, advancement of the gut microbiome composition, enhancement of the antioxidant activities in the colon, liver, kidney, and heart tissues of the host, and decrement of dyslipidemia and blood pressure. However, the type and intensity of the aforementioned effects vary from strain to strain [10-13].

Each strain of these microorganisms that have been recognized as probiotics has unique and unusual properties which vindicates the studies for new probiotics. The biological activities of probiotics are strain-specific and these properties cannot be hypothesized to other strains of the same species. Therefore, it is important to reveal these properties through physiological and metabolic tests, and probiogenomic approaches when an attention-grabbing new strain is recovered from any niche [14-16].

The Shalgam is a traditional lactic acid fermented Turkish beverage that is characterized by reddish color, sour-soft taste, and cloudy appearance [17]. It has been reported that Shalgam exhibits presumptive health benefits against a variety of health risks and diseases due to the high content of lactic acid bacteria present in its microbiome [14, 18, 19]. In its microbial distribution, *Lactiplantibacillus plantarum* and *Lactocaseibacillus paracasei* are reported as prevalent strains by several authors [19, 20]. Except those, *Lb. casei, Lb. brevis, Lb. parabrevis, Lb. pentosus, Lb. buchneri, Lb. delbrueckii* subsp. *delbrueckii, Lb. fermentum, Lb. reuteri, Lb*.*helveticus, Lb. gasseri, Lb. acidophilus, Pediococcus* spp., *Leu. mesenteroides* subsp. *mesenteroides/ dextranicum, Leu. mesenteroides* subsp. *cremoris, Leu. mesenteroides* subsp. *mesenteroides* species were also reported to be found in the shalgam microbiome [18, 21].

This study has aimed to evaluate the genomic, physiological, and metabolic characteristics of *Limosilactobacillus fermentum* strain AGA52 by using “*in vitro*” and “*in silico*” approaches. The strain AGA52 has been isolated from shalgam and as far as we know, there have been no studies regarding the genomic and phenotypic characterization of a *Limosilactobacillus fermentum* strain sourced from traditional lactic acid fermented beverage called “shalgam”, although several kinds of research have been done to enlighten its microbiota, chemical, and volatile composition relationship. For this purpose, the strain’s physiological properties were examined under the screening and selection criteria of probiotics. Further, the whole genome of the AGA52 was sequenced and analyzed with bioinformatic tools. This is the first report clarifying genomic and metabolic characteristics of cholesterol-reducing and GABA-producer *Limosilactobacillus fermentum* strain AGA52 isolated from lactic acid fermented Shalgam beverage.

## Materials and Methods

### Growth Conditions of Bacterial Strain

*Limosilactobacillus fermentum* strain AGA52 was isolated from a Turkish fermented shalgam juice (pH: 3.29) that was purchased from a local producer (Ali Göde: Hot) in Adana, Turkey. A 10 mL of sample was diluted with 90mL of sterile physiological saline solution (0.85%) and homogenized for 1 minute with a highspeed vortex (MS-3 Basic, IKA-Werke GmbH, Staufen, Germany). Later, serial decimal dilutions were get set from the suspension and 100µL of each diluent was spread on MRS agar (Merck GmbH, Darmstadt, Germany). The plates were incubated at 30°C for five days in an anaerobic atmosphere. The AGA52 isolate has been chosen from the dilution of 10^−5^ and subjected to colony purification twice and, catalase test and gram staining were applied to the pure isolate of the AGA 52. The cryo stocks of AGA52 were prepared in MRS broth (Merck) with 25% glycerol and were stored at -80°C.

### DNA extraction, identification, whole-genome sequencing, and assembly

First, *Lb. fermentum* AGA52 cryo culture was subcultured twice in MRS broth (Merck) followed by incubation anaerobically at 37 °C for 24h. A 1 mL fresh culture was pipetted onto a sterile microcentrifuge tube (2mL) and centrifuged at 6000 x g for 10 min at 4 °C. Later, Total DNA was harvested from the cell pellet using the PureLink Genomic DNA Mini Kit (Invitrogen, Thermo-Fisher Scientific, Carlsbad, CA, USA) in accord with the manufacturer’s instructions for gram-positive bacteria. The quality and concentration of genomic DNA were checked by a Qubit 3.0 fluorometer (Invitrogen, Thermo-Fisher Scientific, Carlsbad, CA, USA) and agarose gel (1.5 %). The test strain was determined by full-length nucleotide sequencing of the 16S rRNA gene. PCR amplification and further steps were carried out as per Yetiman et al. [14]. Following the verification of the strain AGA52 using 16S rDNA, the whole genome sequencing libraries were constructed using Nextera XT DNA Library Preparation Kit (Illumina, San Diego, CA, USA) and sequencing was carried out by Illumina Novaseq platform as paired-end (PE) 2x150 bases read. The adapter sequences were trimmed with the JGI-RQC Filter pipeline (BBTools v38.22) by preserving the reads belonging to the strain AGA52 and trimmed raw data were assembled with SPAdes v. 3.15.3 [22].

### Bioinformatic analyses

Genome annotation was performed by using NCBI Prokaryotic Genome Annotation Pipeline (PGAP) [23]. Next, RASTtk and BV-BRC annotations have been conducted for comparison with PGAP [24, 25]. A BLAST Ring Image of the genome of the AGA52 strain and other compared *Lb. fermentum* strains were generated using BRIG v0.95 [26]. The calculation of orthologous average nucleotide identity values (OrthoANI) of the AGA52 and others compared *Lb. fermentum, Lb. gastricus*, and *Lb. reuterii* strains were executed by OrthoANI tool v0.93.1 [27]. Hierarchical clustering analysis of the strain AGA52 and other 45 *Lb. fermentum* genomes were done via the genome clustering service of IMG/M [28]. Prediction of metabolic pathways of *Lb. fermentum* AGA52 was carried out using BlastKOALA for scanning against the KEGG database [29]. Prediction of CAZymes (Carbohydrate active enzymes) was implemented by use of dbCAN2 meta server [30]. A cluster of Orthologous Groups (COG) prediction based on protein FASTA sequences was conducted via webMGA with default options [31]. The upSet plot has been visualized by an intervene web-basedsed intersection tool for the determination of the shared COG between the AGA52 and other reference genomes [32].

The prophage elements on the genome of AGA52 were identified with the PHASTER-Phage Search Tool Enhanced Release [33]. All protein-coding sequences obtained from PHASTER have been screened against the non-redundant protein (NR) database by performing protein-BLAST to identify the horizontally transferred genes. If a gene’s homologous protein was got to match a microorganism other than *Lb. fermentum* ≥ 80%, that gene was noted as horizontally transferred [14]. A resistome screening was conducted by scanning the complete genome sequences of the AGA52 strain versus the ResFinder 4.1, CARD, BV-BRC and KEGG databases, respectively [24, 29, 34, 35]. As with the phage elements, horizontal gene transfer screening was performed within the detected resistome. Prediction of CRISPR-Cas regions were implemented via the CRISPR finder online tool (http://crispr.i2bc.paris-saclay.fr/Server/). The complete genome sequence of *Lb. fermentum* AGA52 was submitted to NCBI under accession number (CP091132.1).

### Carbohydrate fermentation

According to the manufacturer’s protocols, the carbohydrate fermentation patterns of the AGA52 strain were determined by API 50 CHL kit (BioMérieux, Marcy l’Etoile, France).

### Determination of antibiotic susceptibility

Antibiogram assays were performed to find out the resistance or sensitivity of the strain AGA52 in return for commonly used antibiotics. Ready to use commercial antibiotic disks [azithromycin, methicillin, vancomycin, amikacin, kanamycin, tetracycline, penicillin G (Bioanalyse, Yenimahalle, Ankara, Turkey); amoxicillin, amoxicillin, ampicillin, erythromycin, oxacillin, carbenicillin, streptomycin, rifampicin (Oxoid, Basingstoke, Hampshire, UK)] were used for antibiotic susceptibility testing of *Lb. fermentum* AGA52. The application of disk diffusion assay was carried out considering the Kirby-Bauer method [36] Interpretation of inhibition zone (mm) was carried out as described by Clinical and Laboratory Standards Institute’s performance standards for antimicrobial testing [37].

### Determination of probiotic properties

To determine probiotic properties of the AGA52: β-haemolysis, *in vitro* simulation of gastric digestion, cell surface hydrophobicity, cellular auto-aggregation tests, and antibacterial activity assay against several pathogens were performed, respectively. β-hemolytic activity of the AGA52 was evaluated by using 5% sheep blood containing Columbia agar plate. The isolate was streaked on the Columbia agar followed by incubation at 37°C for 48 h under anaerobic conditions [38]. To check the response of the AGA52 against gastric digestion, the gastric environment was simulated as described by Zhang et al. [39] with a slight modification (1% bile salt was used) for preparation of artificial gastroenteric juice [14]. Cell surface hydrophobicity and auto-aggregation assays were applied as for that Krausova et al. [40]. The bactericidal activity of the strain AGA52 has been tested by using the agar well diffusion method [41]. The supernatant of 48h grown bacterial cells was assayed against *Escherichia coli* O157:H7 (ATCC 43895), *Staphylococcus aureus* (ATCC 25923), *Bacillus cereus* (ATCC 33019), *Salmonella enterica* sv. *Typhimurium* (ATCC 14028), *Proteus vulgaris* (ATCC 8427), *Listeria monocytogenes* (ATCC 7644), *Yersinia enterocolitica* (ATCC 9610) and *Klebsiella pneumoniae* (ATCC 13883).

### Antioxidant Activity Assays

The antioxidant activity of the AGA52 was determined by using DDPH (2,2-diphenyl-1-picrylhydrazyl) and ABTS (2,2 azino-bis(3-ethylbenzothiazoline-6-sulfonic acid)) methods. For this purpose, cell-free supernatant (CFS) was collected from MRS broth via centrifugation at 3500rpm for 15 min. DPPH scavenging activity was assessed according to Ozturk et al. [42] with some modifications. The reaction mixture was prepared by adding 2 mL freshly prepared 0.2mM DPPH onto 1 mL four times diluted CFS. The mixture was incubated in the dark for 30 min at 25°C. Later, the mixture was centrifuged at 5000 rpm before measuring the absorbance at 520 nm with five replicates. The blank control includes dH_2_O and DPPH solution. The DPPH scavenging activity has been computed as scavenging activity % = (1 − A_test_/A_blank_) × 100.

ABTS assay was performed as for that Pieniz et al. [43] with some modifications. ABTS was dissolved in water (7 mM). Later, ABTS stock solution was mixed with 2.45 mM potassium persulfate (final concentration) for the formation of radical cation (ABTS^+^) and the mixture was incubated in dark for 16h at 25°C before usage. The stock solution can be used for only 3 days. Before application, ABTS^+^ solution was diluted with phosphate buffer (pH: 7.2) to get an absorbance of 0.700±0.03 at 734 nm. Likewise, The CFS was also diluted four times with phosphate buffer (pH: 7.2). Next, 40 mL of CFS was added to 4mL of ABTS^+^ solution, the mixture was incubated in dark for 5 min and the absorbance was measured with five replicates. Phosphate buffer (pH: 7.2) was used as blank and control. The percentage of inhibition was predicted according to the ascorbic acid standard curve (0-9 µg/mL).

### Cholesterol assimilation assay

The cholesterol utilization by the AGA52 in MRS broth was analysed by O-phthaldehyde (OPA) method as explained by Rudel, Morris [44] with some modifications. The AGA52 was grown in MRS broth (0.25% dextrose + 0.3% ox bile containing) supplemented with 100ppm cholesterol (5-cholesten-3β-ol (Sigma, Merck GmbH, Darmstadt, Germany), dissolved in 2-propanol) at 37 °C for 24 and 48h, respectively. After incubation, the cultures were centrifuged (3500rpm, 15min, 25°C) and CFS were used for determination of residual cholesterol. Next, 0.5mL of CFS was mixed with 2mL of KOH (50% wt/vol) and 3mL of absolute ethanol, vortexed for 1 min and later heated at 60°C for 15 min. After chilling, 3mL dH_2_O and 5mL of hexane were added to the mixture and it vortexed for 1 min. Afterwards, 2.5 mL of hexane phase was transferred to another glass tube and then was kept for evaporation at 80°C. The residue was directly dissolved in 4 mL of OPA (0.5 mg/mL in glacial acetic acid) and incubated at 25°C for 10 min. Later, 2 mL of H_2_SO_4_ (98%) was added gently and vortexed for 1min and the mixtures incubated again at 25°C for 10min, and absorbance was measured at 550nm (Shimadzu UV-1800 UV/VIS spectrophotometer, Tokyo, Japan). The cholesterol assimilation was calculated according to the difference between the control (uninoculated MRS broth) and test samples.

### Confirmation of GABA production by ^1^H-nuclear magnetic resonance

The monosodium-glutamate (MSG) standard, γ-aminobutyric acid (GABA) standard and deuterium oxide were purchased from Merck GmbH (Darmstadt, Germany). The two-times subcultured AGA52 culture was inoculated in 1% MSG containing MRS broth (0.25% dextrose) and incubated at 37 °C for 48h. Next, the culture media was centrifuged at 3500 rpm for 15 min and cells were discarded. The CFS was lyophilised by using a freeze-dryer (Christ, Alpha 2–4 LSCplus, Germany). Next, aprroximately 100mg of CFS, MSG, and GABA were transferred into a cylindiric glass NMR tube and dissolved with deuterium oxide, respectively. Each sample was subjected to ^1^H-nuclear magnetic resonance by using an NMR spectrometer (Bruker, Model 400, MA, USA) at 400mhz for 5 min. Afterwards, NMR spectra were analysed by comparing standards via data analysis software (Bruker, TopSpin, MA, USA) of the device.

## Results and Discussion

### Functional genomic characterization

The genome of the *L. fermentum* strain AGA52 comprised a circular chromosome of 2,001,184 bp with a 51.82% GC ratio, and it contains 2.024 genes, 1.873 protein-coding sequences, 50 tRNA, 3 rRNA, 3 non-coding RNA, 1 CRISPR array region and 95 Pseudogenes. The BLAST atlas of the genome of AGA52 and other compared well-known *L. fermentum* genomes were displayed in Fig 1, and differences between each other can be noticed clearly due to the gaps in the genomes. Among these gaps, the detection of genomic islands is possible due to the corresponding decrease in GC content and the absence of these islands in other strains. Generally, integrases, insertion sequences, and transposases are positioned in and/or around these islands which is an indicator of potentially horizontal transferred genes [45, 46].

**Fig 1.**
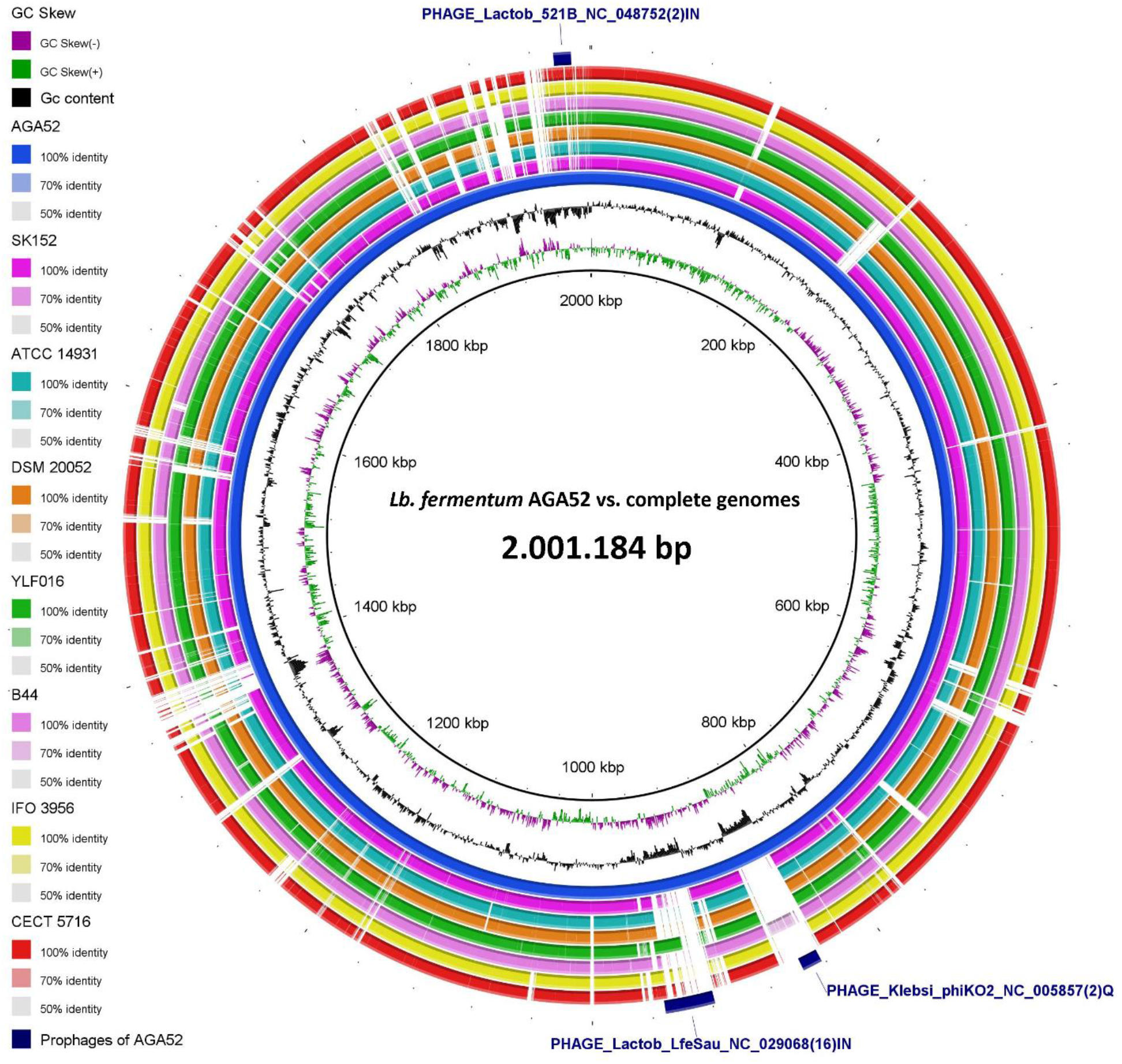
The BLAST Ring alignment of *Limosilactobacillus fermentum* strain AGA52 versus other compared well-known *Lb. fermentum* genomes which were aligned from the red ring to the inside as CECT 5716, IFO3956, B44, YLF016, DSM 20052, ATCC 14931, SK152, and AGA52, respectively. The GC content was illustrated in the third inner circle before the genome of AGA52. The GC skew (+/-) of the genome of AGA52 was also shown in the 2nd inner circle and in the first inner circle, the genome size has been demonstrated. On the other hand, prophage regions of the strain AGA52 were depicted as navy blue colored arcs in the outer circle, the numbers in parentheses show the number of matched proteins of the phages. Besides, at the end of the phage names, (IN) means intact and (Q) means questionable.

The orthoANI comparison results amongst the AGA52 and the aforecited genomes were presented in Fig 2. Based on the OrthoANI values, the genome of the *L. fermentum* AGA52 bears a resemblance to the genomes of SK152 (99.18%), ATCC 14931 (99.15%), DSM20052 (99.11%), YLF016 (99.11%), IFO 3956 (99.04%), CECT 5716 (98.92%), and B44(98.92%), respectively. It has been informed that the recognition of the genomes of the two species as the same depends on the ANI value above 96% [27]. Besides, It has been known that SK152, ATCC14931 and DSM 20052 originated from fermented plant materials [45, 47]. On the other side, YLF016 (Yak’s GIT), CECT5716 (human breast milk), and IFO3956 (cheese) are derived from non-plant sources [5, 39, 48]. The genetic distance between the genome of AGA52 and the mentioned genomes can be explained by differences or similarities in the strain’s isolation niche, evolutionary histories, and the conditions exposed [14, 49]. In addition, assembly details and annotated genome properties of *L. fermentum* AGA52 and higher orthoANI similarity shown genomes were summarized in Table 1.

**Table 1.** Summary of assembly details and annotated genome properties of 986 *Limosilactobacillus fermentum* AGA52 and other compared strains which have higher 987 orthoANI similarity to itself.

| Item | AGA52 | SK152 | ATCC 14931 (B1 28) | YLF016 |
| --- | --- | --- | --- | --- |
| Contigs | 128 | 1 | 1 | 1 |
| Genome length (bp) | 2,001,084 | 2,092,273 | 1,905,587 | 2,094,354 |
| GC content (%) | 51,82 | 51,94 | 52,3 | 51,46 |
| Sequencing platform | Illumina<br>Novaseq | PacBio | Oxford Nanopore<br>GridION | PacBio |
| Sequencing depth | 296.0x | 500.0x | 342x | 200.0x |
| Genes (total) | 2,024 | 2,121 | 1,916 | 2,136 |
| Protein coding sequences | 1,873 | 1,950 | 1,744 | 1,950 |
| tRNA | 50 | 58 | 58 | 58 |
| rRNA | 3 | 15 | 15 | 15 |
| Non-coding RNA | 3 | 3 | 3 | 3 |
| CRISPR_repeat | 15 | 100 | 13 | 10 |
| CRISPR_spacer | 14 | 98 | 12 | 9 |
| CRISPR_array | 1 | 2 | 1 | 1 |
| Pseudogenes (total) | 95 | 95 | 96 | 110 |
| Pseudogeens (frameshifted) | 19 of 95 | 28 of 95 | 36 of 96 | 35 of 110 |
| Pseudogenes (incomplete) | 68 of 95 | 50 of 95 | 35 of 96 | 60 of 110 |
| Pseudogenes (internal stop) | 24 of 95 | 29 of 95 | 33 of 96 | 30 of 110 |
| OrthoANI similarity to AGA52 | - | 99.18 | 99.15 | 99.11 |

**Fig 2.**
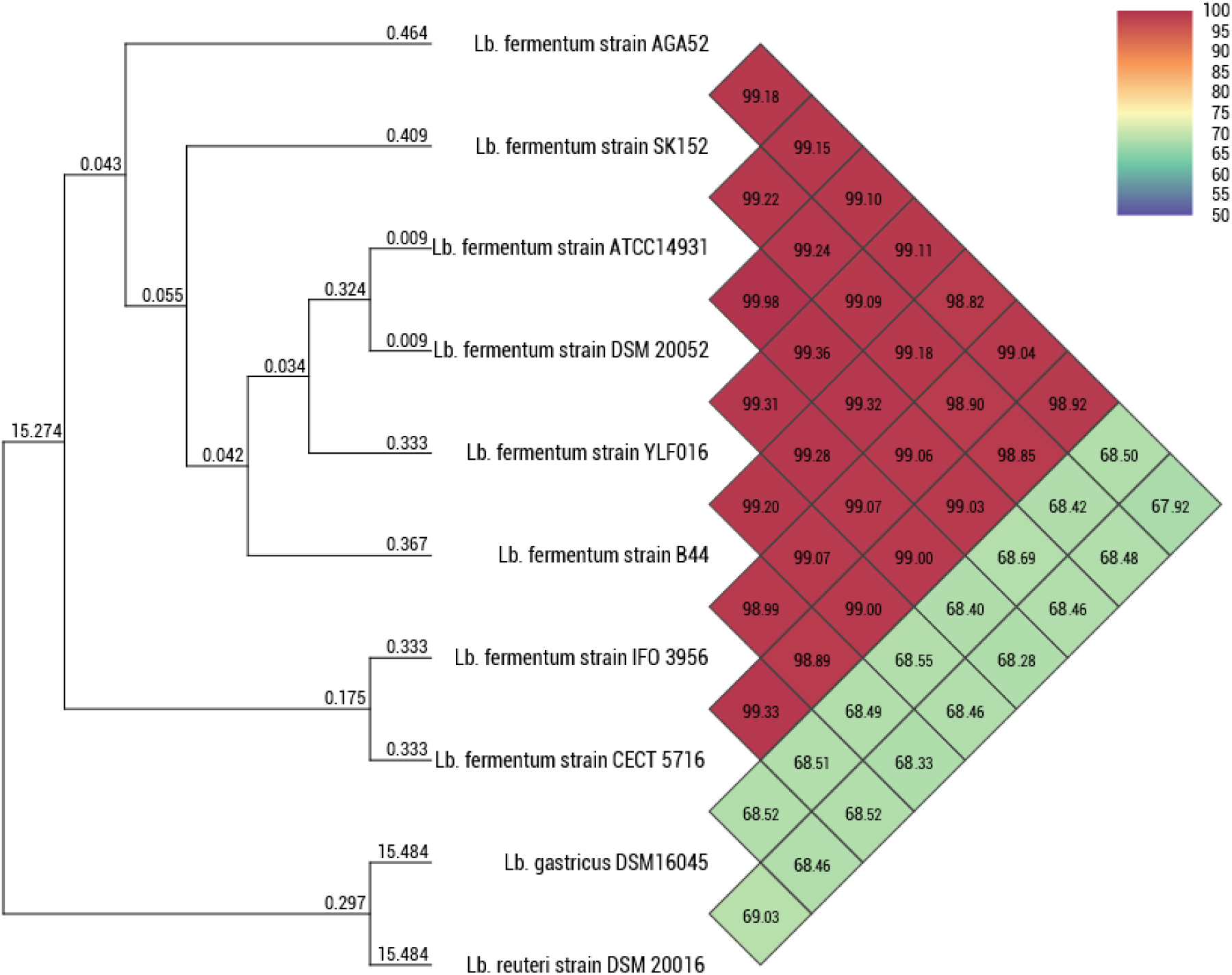
Orthologous Average Nucleotide Identity (OrthoANI) phylogram of the genomes of the *Limosilactobacillus fermentum* strain AGA52 and other well-known *Limosilactobacillus* strains.

Moreover, the protein-coding sequences of the genome of the AGA52 have been screened for determination of the clusters of orthologous groups (COG) and categorized into 23 different function classes in Fig 3. A detailed demonstration of the number of shared orthologous proteins by the UpSet plot can be seen in Fig 4. According to the hierarchical clustering results (Fig 5.) which are based on clusters of orthologous groups (COG) profiles of 46 different *L. fermentum* strains, the AGA52 shared the highest similarity with KMB612 (0.95), KMB613 (0.95), AF15-40LB (0.95), AF16-22LB (0.95), 279 (0.94), 779_LFER (0.94), and VRI-003 (0.94). The isolates of KMB 612 and KMB 613 were isolated from the Bryndza cheese (Fig S1). The characteristic of cheese is that it is produced from sheep’s milk. Based on past studies on *L. fermentum*, It is possible to make certain assumptions to explain this genetic relationship between AGA52, KMB 612, and KMB 613. This strain may have been transferred to sheep’s milk through the consumption of plant-based fermented feed (e.g., silage) or may itself be an autochthonous member of the existing sheep microbiome [45, 49]. In the IMG/M database, AF15-40LB, AF16-22LB and 279 genomes were reported as faecal originating, while 779_LFER has been expressed to originate from the respiratory system. It was also described in the IMG/M that VRI-003 was isolated from a lyophilized commercial probiotic formulation from Australia. Nonetheless, It has been reported that VRI-003 has an increasing effect on interferon-gamma (IFNγ) production in whole blood culture [50]. As above, a similar analogy can be drawn between AGA52 and these strains.

**Fig 3.**
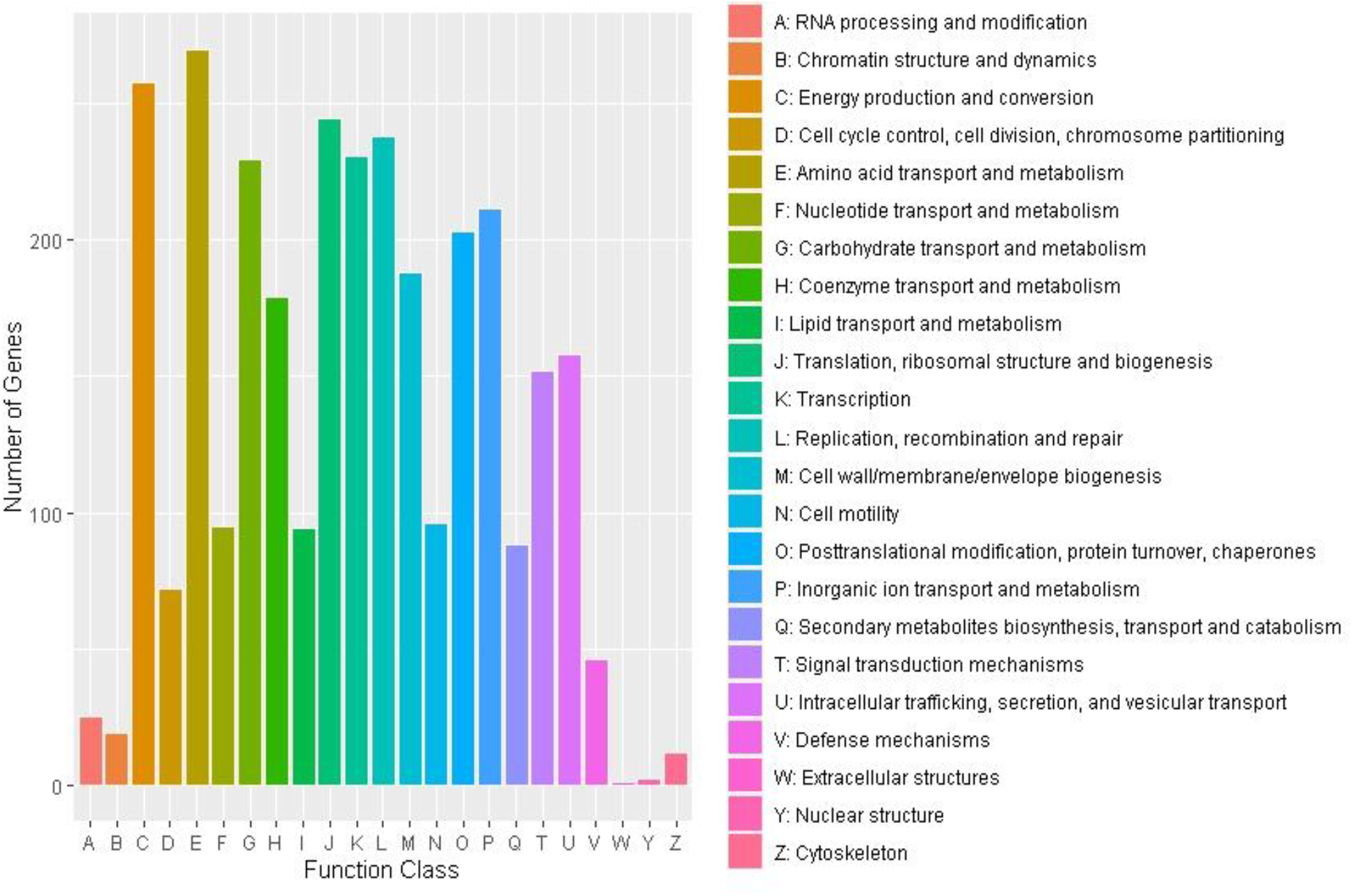
Graphical presentation of the identified numbers of clusters of orthologous groups (COG) of protein-coding sequences in the genome of *Limosilactobacillus fermentum* AGA52.

**Fig 4.**
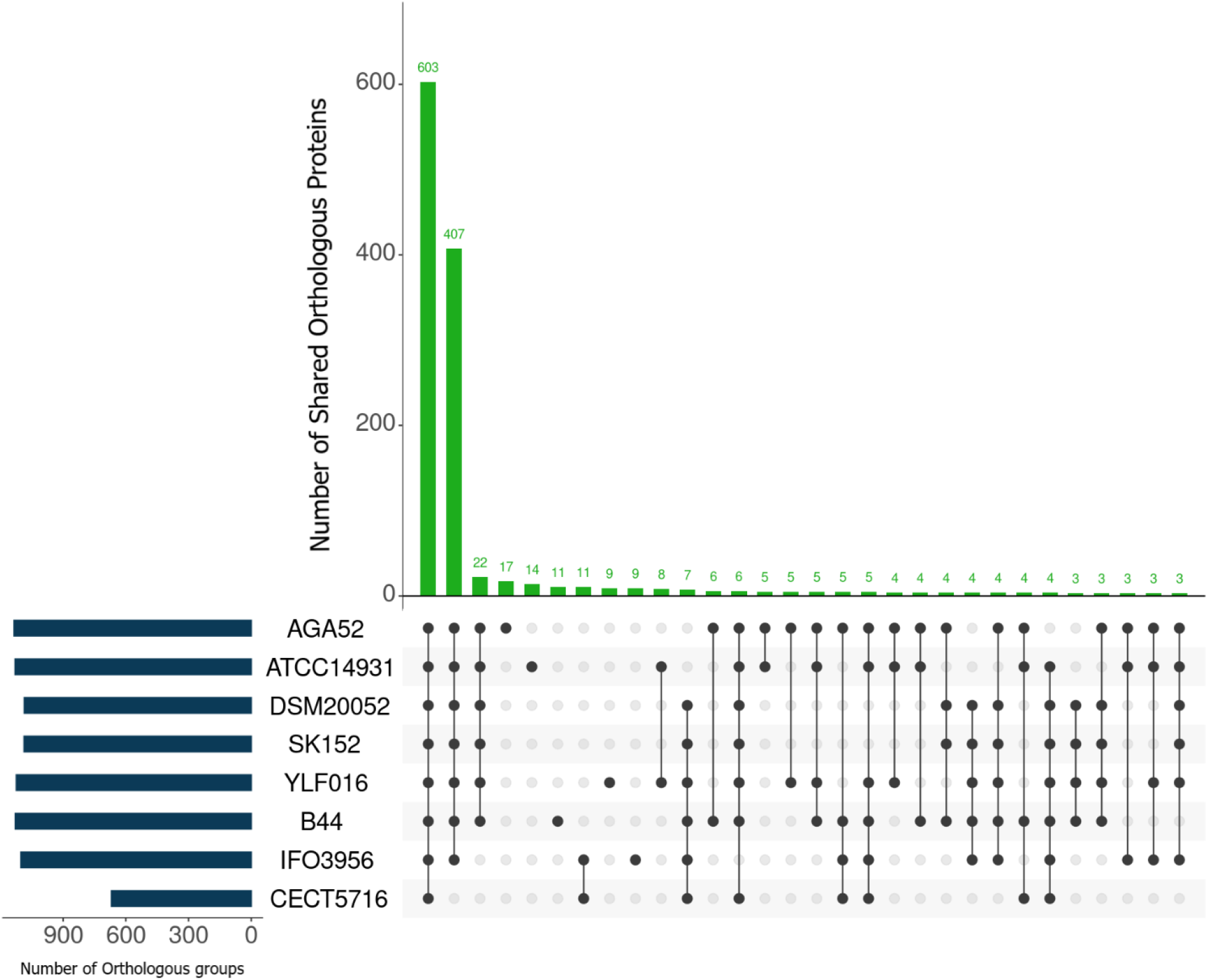
The UpSet plot demonstrates the number of shared orthologous proteins amongst AGA52 and the other well-known strains with bar charts. On the other hand, the number of total identified clusters of orthologous groups (COG) via the webMGA engine has been shown on the left of the strain names.

**Fig 5.**
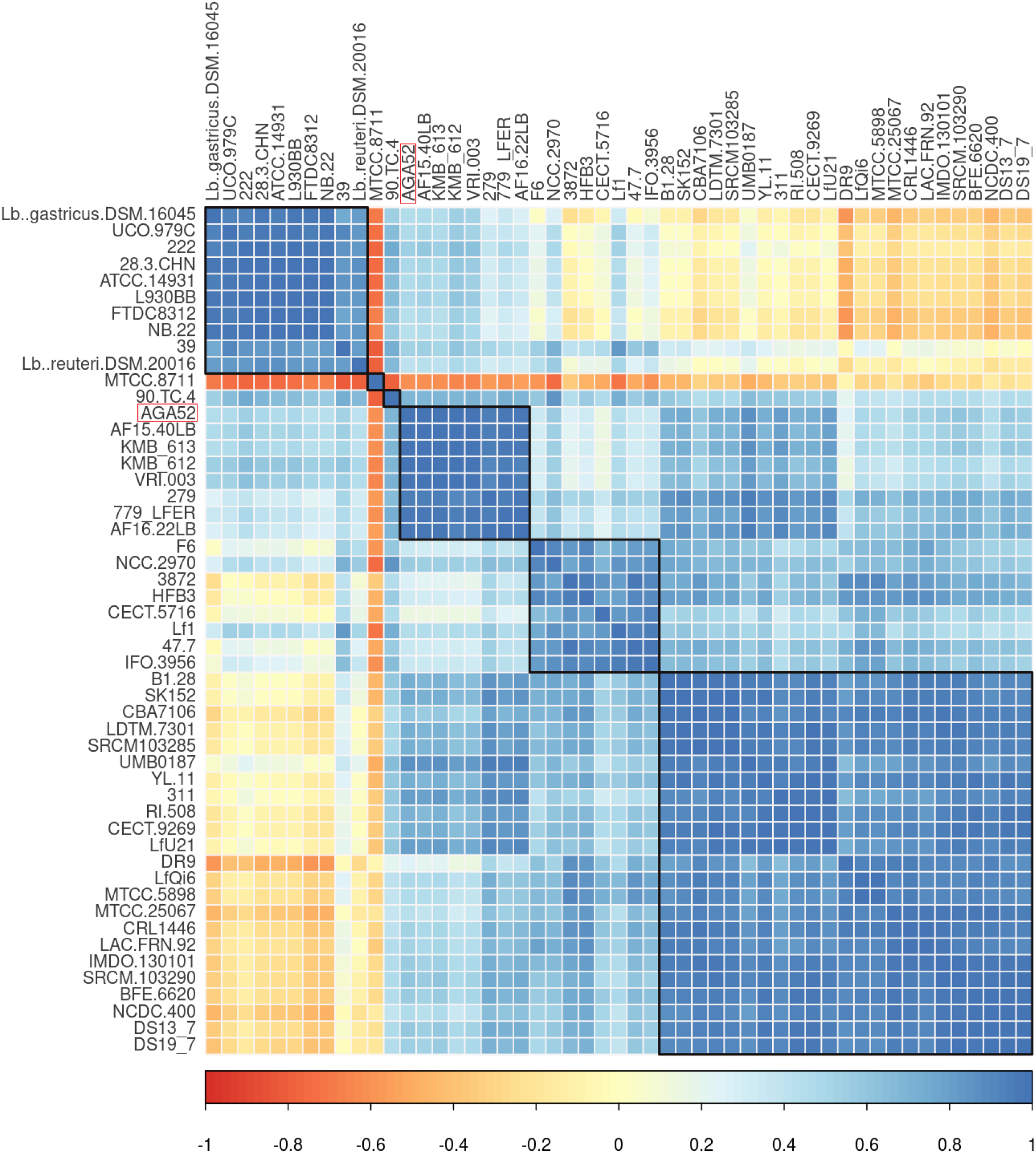
Pairwise intersection with hierarchical clustering in accordance with the existence of clusters of orthologous groups (COG) of 48 different *Lb. fermentum, Lb. reuteri* DSM 20016, and *Lb. gastricus* DSM 16045 genomes which were available on IMG/M. The correlation coefficient was calculated according to Pearson correlation and the color bar on the bottom of the heatmap indicates the degree of the correlation coefficient. The AGA52 shows a higher correlation with KMB612 (0.95), KMB613 (0.95), AF15-40LB (0.95), AF16-22LB (0.95), 279 (0.94), 779_LFER (0.94), and VRI-003 (0.94), respectively. Interestingly, it is apprently seen that MTCC 8711 inversely correlated with the all genomes clustered.

Predicted CRISPR structures in the genome of the AGA52 consist of 1 copy of CRISPR-associated endonuclease Cas9, 1 copy of type II CRISPR-associated endonuclease Cas1, 1 copy of type II-A CRISPR-associated protein Csn2, 1 copy of CRISPR-associated endonuclease Cas2, 15 repeat regions, 14 spacer regions, and 1 array region (Table S1). CRISPR constructs and related (Cas) proteins serve as adaptive immune systems against prophages and foreign plasmids in prokaryotes and have been used for genotyping and genetic engineering purposes since the day they were discovered [51, 52]. Especially, from the point of view of the development of next-generation probiotics with enhanced functionalities, this technology has uncovered new opportunities. In this sense, It is possible to improve the therapeutic potential of the probiotics for vaccine delivery or modulation of the host immune response [53]. Therefore, the prediction and characterization of the CRISPR-Cas structures in the genome of the *L. fermentum* strain AGA52 could enable the development of novel biotherapeutics by regulating the expression and production of existing functional metabolites or acquiring novel biosynthetic gene clusters (e.g bacteriocins, non-bacteriocin antimicrobial compounds, metabolic products) by using itself.

### Prophages, related horizontal gene transfer, and mobilome

The phage scan results revealed that AGA52 had three phage sites in its genome, two intact and the other questionable. The intact phage regions (regions 2 and 3) were displayed similarity with Lactob_LfeSau_NC_029068 (32.2 kb) and Lactob_521B_NC_048752 (11.7 kb), respectively. It has been observed that Lactob_LfeSau presented the highest matching protein (16) among the identified phages which is also shown in Table S2. This phage is the most common one found to be among the *L. fermentum* strains. Only, Lactob_LfeSau has attL/attR sequences and integrase (PP_00944) except for region 1 (Klebsi_phiKO2) and 3 (Lactob_521B). In a bacterial genome, the presence of the integrase is a functional identifier for pathogenicity islands, phages, and integrative plasmids [46, 54]. Besides, GC% content of the phages is quite lower than the average GC% content of the genome of the AGA52. All phage regions have been located around or in the genomic islands which are seen in BLAST ring alignment (Fig 1). Both attachment sites are positioned forward strand and upstream of the integrase (attL: 922221-922233; attR:947491-947503). Merely, Lactob_LfeSau was found to comprise all packaging/head/tail gene clusters, and DNA packaging genes. Unlike, previous studies no endolysin gene has been observed among all of the identified phages [55, 56]. Endolysins are phage proteins that rapidly break the bacterial cell wall down and release new viral particles [57]. This means that identified phages are temperate for the strain AGA52 and each member of identified phage regions was listed in Table S3-S5.

Horizontal gene transfer (HGT) amidst bacteria generally appears in so far as bacteriophage infection or natural competence [14, 58]. Based on protein BLAST results for screening HGT from phage-related proteins, 0.79% (16 genes) of the genome of AGA52 have been acquired by phage-associated HGT from other bacteria and viruses, eg. *Ligilactobacillus animalis, Pediococcus acidilactici, Lactobacillus* sp., *Siphoviridae* sp. (bacteriophage), *Lentilactobacillus* sp., *Limosilactobacillus reuteri* (Table S6). On the other side, all of the detected phage-associated transposases were only found in the Lactob_521B phage. Among the phage elements of Lactob_521B, 11 transposase genes (PP_01962, PP_01963, PP_01965, PP_01966, PP_01968, PP_01970, PP_01972, PP_01973, PP_01975, PP_01976, and PP_01977) that may have been acquired by recombination, repair and replication. In addition, as for the whole genome-based transposase search result, IS3, IS4, IS6, ISL3, IS30, IS200/IS605, IS256, and IS982 families’ members of transposases have been identified by using IS Finder and were summarised in Table S7.

### Phenotypic antibiotic resistance and Safety-related genes assessment

In agreement with the European Food Safety Authority (EFSA) suggestions, a probiotic strain should not hold gained or transferable resistome to clinically used drugs [59]. Therefore, it is important to determine antibiotic resistance phenotypically and genotypically. Assessment of the antibiotic susceptibility of the AGA52 has been performed following The Clinical and Laboratory Standards Institute’s performance standards. The value of inhibition zones (ZOI) that belongs to tested antibiotics against the AGA52 and resistome search results were presented in Table S8. Based on the antibiogram results, AGA52 was found to be sensitive (ZOI ≥ 20mm) to Ampicillin (10 μg), Penicillin G (10 U), Carbenicillin (100 μg), Amoxycillin (25 μg), and Rifampicin (5 μg), while it displayed resistance (ZOI ≤14mm) to Methicillin (5 μg), Oxacillin (1 μg), Streptomycin (10 μg), Vancomycin (30 μg), Amikacin (30 μg), and Kanamycin (30 μg). Unlike these, intermediate sensitivity (ZOI ∼ 15-19mm) to Azithromycin (15 μg) and Erythromycin (10 μg) was also observed.

Based on Resfinder and CARD analyses, there are no antibiotic-resistance genes have been found in the genome of the AGA52. Nonetheless, sixteen antibiotic resistance genes have been found by using KEGG and BV-BRC databases which were displayed in Table S8. The detected genes were found to be related to β-Lactams (4), aminoglycosides (7), peptide-antibiotics (3) and multidrug efflux transporters (2). It is well known that lactobacilli show a higher resistance profile to aminoglycosides. Especially, *Lactobacillus* species have been acknowledged as resistant to vancomycin under their intrinsic peptidoglycan precursors containing D-lactate instead of D-alanine at the C-terminus [60]. The vancomycin resistance genes were detected in the genome of the AGA52 comprises of *mraY, alr, ddl, murF, murG*, and *VanS* which were presented in Table S9. Interestingly, detected vancomycin resistance genes showed differences from previous studies [14, 39]. Especially, the *VanX* gene could not be found which is highly specific for hydrolyzation of D-ala-D-ala dipeptides and is an important precursor of the cell wall. This might be caused by variations in tolerance or gene expression [61]. Additionally, as an aminoglycoside, streptomycin resistance responsible *gidB* gene has also been observed. However, no specific resistance genes were identified for kanamycin and amikacin even if resistance subsists. This could be caused by phenotype do not completely represent genotype [62]. Aside from these, *blaI, mcrA, pbp2A*, and *penP* genes responsible for β-Lactam resistance have been determined which can be seen in Table S8. It is well-known that oxacillin and methicillin resistance is related to penicillin-binding proteins (*mcrA, pbp2A*). Despite the presence of beta-lactamase (*penP*) and penicillinase repressor (*blaI*), AGA52 exhibited a sensitive profile to ampicillin, penicillin, carbenicillin, and amoxycillin. This would be explained by the aforementioned case for aminoglycosides. On the other hand, there are no resistance genes were found for the macrolides, tetracyclines, and rifamycins tested in this study. However, strain AGA52 exhibited an intermediate and sensitive pattern to these antibiotics (Table S8). This situation could be derived from membrane impermeability and/or multidrug efflux transporters (the presence of *efrA* and *efrB* is confirmed in Table S9 and S12) in reduced susceptibility to these drugs [63]. In addition, daptomycin-specific *pgsA, mprF*, and *gdpD* genes were also identified in the genome of the AGA52. Due to increasing concerns regarding various food niches and/or common bacteria that might serve as potential reservoirs for resistome, the presence of transferable antimicrobial resistance genes in probiotics is undesirable which is used for humans or animals. Lactobacilli have intrinsically originated resistance to various antimicrobial compounds, and it is well-known that such resistance has not been related to any particular safety concerns. But, the intrinsic resistome on the bacterial chromosome must not be flanked by transposases or integrases. According to the protein BLAST results for the resistome detected in this study, no horizontally transferred gene was found (Table S9).

### Carbohydrate fermentation patterns and active enzymes

*Limosilactobacillus fermentum* acquires ATP through heterofermentative carbohydrate fermentation based on strain-specific sugar preference. Uncovering strain-specific metabolic tendencies is important due to the prediction of industrial applicability and their probiotic potential. Since, carbohydrate-active enzymes (CAZYmes) also have a probiotic function, e.g. xylose-isomerase stimulation of gut persistence [64]. According to API 50 CHL test results, the AGA52 can utilize 13 different carbohydrates out of 49 being tested. At first glance, the utilization of D-ribose, D-xylose, D-galactose, D-glucose, D-fructose, D-cellobiose, D-maltose, D-lactose, D-melibiose, D-sucrose, and D-raffinose can be comprehended from Table S10. *L. fermentum* is a cohesive species and could ferment various kinds of sugars. This status is directly involved in the presence of the genes responsible for the CAZYmes and transporters. Most of the transporters associated with carbohydrate metabolism are positioned in the phosphoenolpyruvate-dependent sugar phosphotransferase system (PTS) [65]. For strain AGA52, the entire PTS enzymes were encoded by its genome and it consists of PTS System Enzyme I (general enzyme gene, *ptsI*, K08483), phosphocarrier protein HPr gene (*ptsH*, K02784), and 13 complete or incomplete substrate-specific enzyme II (EII) complexes genes (Table S13). These complex genes seem cellobiose, fructose, mannose, sucrose, beta-glucoside, and ascorbate specific and their substrate specificity is unidentified. However, it is well-known that various sugar transporters can bring in more than one substrate [58]. In addition, several carbohydrate-specific common ABC transporters were listed in Table S12.

It has been observed that the AGA52 has not encoded 1-phosphofructokinase (*pfK*) enzyme gene on its genome. Based on previous studies, it is commonly accepted that species lacking this enzyme have an obligate heterofermentative carbohydrate metabolism and they also produce CO_2_, lactate, and ethanol via a heterofermentative pathway [66, 67]. The same case is valid for AGA52. Moreover, the presence of l-arabinose isomerase, l-ribulose kinase, and ribulose phosphate epimerase enzymes are confirmatory to a lesser extent that a lactobacilli strain has an obligate heterofermentative pathway [2, 68]. In this study, both L-arabinose isomerase [EC:5.3.1.4] and ribulose-phosphate 3-epimerase [EC:5.1.3.1] could only be found in the genome of the AGA52 (Table S11). Nevertheless, it could be implied that AGA52 has an obligate heterofermentative pathway. Meanwhile, the key enzymes responsible for the EMP, PK and Leloir pathways and their copy numbers are presented in Table S14 by being compared with well-known *L*.*fermentum* strains. Among the compared strains, all key enzymes are available for the aforecited pathways apart from phosphofructokinase 1, fructose-bisphosphate aldolase, and L-ribulose kinase. These findings are also confirmatory for the mentioned situation above. The CAZYmes encoding genes connected with the intact EMP (Fig S1) and PK (Fig S2) pathways have been identified in the genome of the AGA52 and presented in Table S11.

The AGA52 can synthesize both isomers of lactate and has got four copies of L-lactate dehydrogenase (*ldh*), three copies of D-lactate dehydrogenase (*ldhA*), and one copy of D-lactate dehydrogenase (*dld*; quinone) enzymes encoding genes on its genome. The difference between *dld* and others is that it is a peripheral membrane-specific dehydrogenase which plays a role in respiration. The electrons originating from D-lactate oxidation are turned over to membrane soluble quinone pool [69]. Apart from these, the AGA52 also possesses two copies of alcohol dehydrogenase (*adhP*) enzyme encoding genes. The reaction products of this enzyme differ depending on the substrate catalyzed. The enzyme produces aldehydes as a product in reactions where primary alcohols are substrates, while ketones are released as products when secondary alcohols are substrates [70]. The effect of the reaction products on the specific flavour of the shalgam cannot be ignored.

Following KEGG mapper results, the AGA52 has a total of 158 carbohydrate metabolism-related genes which are encoded in its genome. The distribution of those genes is as follows; 18 Embden–Meyerhof–Parnas pathway genes, 14 phosphoketolase pathway genes, 6 TCA cycle-related genes (Fig S3), 6 pentose and glucuronate interconversions associated genes, 13 galactose metabolism genes, 11 fructose and mannose metabolism genes, 3 ascorbate and aldarate metabolism genes, 13 starch and sucrose metabolism genes, 19 amino sugar and nucleotide sugar metabolism genes, 20 pyruvate metabolism genes, 6 glyoxylate and dicarboxylate metabolism genes, 9 propanoate metabolism genes, 13 butanoate metabolism genes, 3 inositol phosphate metabolism genes, and 5 C5-branched dibasic acid metabolism genes. Moreover, the number of identified CAZYmes encoded in the genome of AGA52 is as follows; 32 Glycosyl transferases, 1 auxiliary activity responsible enzyme, 3 carbohydrate-binding modules, and 38 Glycoside Hydrolases. There is no existence of carbohydrate esterases has been observed in CAZY annotation results. Further, it can be seen in Fig 6 that the CAZYmes patterns of ATCC 14931, DSM 20052 and SK152 are similar to AGA52. Overall, the characterization results revealed that the AGA52 has an acceptable potential for carbohydrate degradation and adaptive nature specific to the gastrointestinal tract.

**Fig 6.**
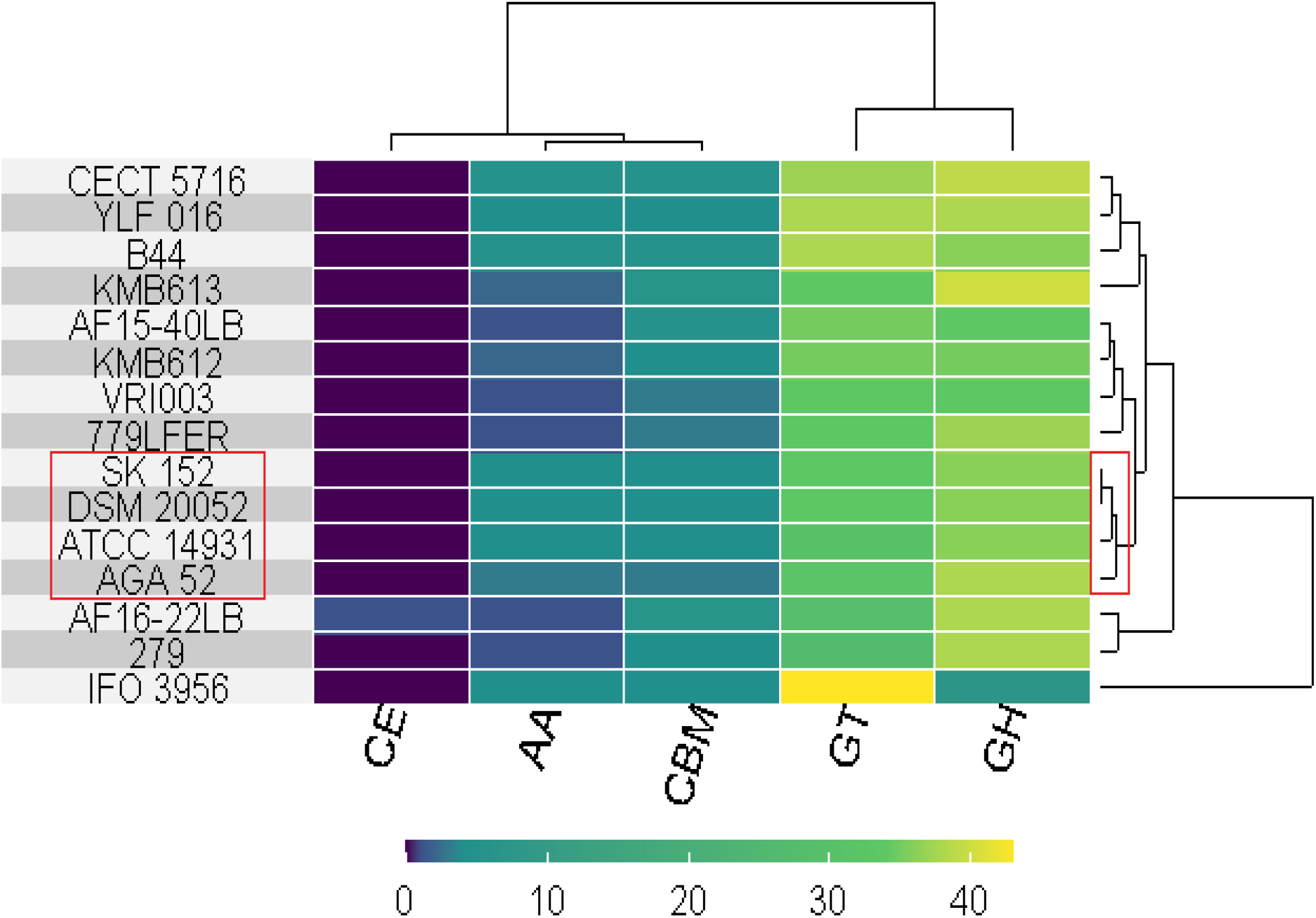
Heatmap of carbohydrate-active enzymes (CAZYmes) and their distribution amongst the compared *L. fermentum* strains. The color gradient bar indicates the exact number of identified enzymes from the CAZY (http://www.cazy.org/) database by using DIAMOND (E-value <1e-102). CE: Carbohydrate Esterases, AA: Auxiliary Activities, CBM: Carbohydrate-Binding Modules, GT: Glycosyl Transferases, GH: Glycoside Hydrolases.

### Probiotic properties and antioxidant activities

The probiotic characterization tests were carried out for confirmation of the presence or absence of the features responsible for becoming probiotics. First, the result of the β-haemolysis assay revealed that AGA52 does not have β-hemolytic activity. The cell surface hydrophobicity (CSH) and autoaggregation capacity (AC) were determined as 73,6% (with xylene) and 44%, respectively. The CSH is essential for understanding probiotics’ overall adhesion capacity, which is generally measured by assessing the bacterial affinity to a hydrocarbon (e.g. xylene, n-hexadecane, hexane) solvent [40, 71]. The hydrocarbon affinity method does not reveal exact CSH, instead, it expresses the presence of van der Waals and electrostatic interactions which are significant factors influencing the overall adhesion capacity [72]. Recently, it has been reported that bacteria which has a higher CSH can bind better to intestinal epithelial cells [73]. Similar to the present study, approximate CSH values were reported for *L*.*fermentum* SJRP40 (79.69%), SJRP46 (80.57%), and *L. reuteri* K18 (76%) by different studies [40, 73]. CSH value of *L. fermentum* YLF016 (≥65%) has been reported as low as than AGA52. Moreover, the lipoteichoic acid (LTA) production responsible genes (*MurJ, LtaS, LtaS, LafC, LafB, LafA, dltD, dltA*, and *dltB*) that can be associated with CSH were identified and their putative functions were explained in Table S15.

Autoaggregation ability for probiotic bacteria is crucial due to its direct relationship to the adhesion of epithelial cells and mucosal surfaces and is also a phenotypic marker for adhesion [74]. Due to the AC, bacteria can generate cellular aggregates which promote persistence in the intestine [75]. Bacterial cell wall components and other structures, like mucus-binding proteins, adhesins, surface layer proteins, fibronectin-binding proteins, exopolysaccharides, and lipoteichoic acids give an advantage to bacteria for colonization and adhesion to host epithelial cells [74, 76, 77]. In this study, the presence of some of these structures has been confirmed in the genome of AGA52 and summarised in Table S15. The AC of the AGA52 shows a similar pattern to previously studied strains such as colostrum-originated *L. reuteri* K14 (44.4%) and infant feces-originated *L. reuteri* E1M1C (46.3%). Besides, Li et al. [78] reported between 0.86 to 65.15% autoaggregation values for a variety of *L. fermentum* strains which were isolated from different Chinese fermented foods.

Resistance to stress factors of gastrointestinal systems, such as long-term survival and colonization are uniquity of Lactobacilli. Uncovering host-probiotic interaction regulation mechanisms have become more significant due to the gradually increasing number of microbes used as probiotics. The survival behaviour of AGA52 was tested against physiological pH, and bile salt environment and mimicked the gastrointestinal fluid. Broadly, survivability at low pH of most Lactobacilli differs between 2.5 to 3.5 which is a crucial selection criterion for potential acid-tolerant probiotic strains. During gastric transit, probiotics must survive at pH 3.0 as low and remain alive for at least four 4 hours before reaching the lower tract [79]. It has been observed that the AGA52 has a tolerance to an acidic environment with a survival rate of 20±0.3% in pH 3.0 (with an initial load of 8.74 log CFU/mL). Besides, the viability goes down to 0.09±0.01% at pH 2.0 (Fig 7B) which overlaps the previous studies [14, 39]. Furthermore, the survival profile to bile salts (0.2-1% (w/v)) was checked and the AGA52 presented a moderate survival pattern (38±0.1%) after 4h (with an initial load of 9.19 log CFU/mL). Commonly, it is strongly suggested that approximately 0.3% bile salts concentration is proper for probiotic selection purposes [80]. Based on the survival pattern at 0.3% bile salt concentration (64±0.1%), the AGA52 appears to partially meet the requirements of this criterion (Fig 7A). Apart from these, survivability of the AGA52 in the gastrointestinal simulated environment was determined for understanding whether the strain shows tolerance or not. Because, gastrointestinal tolerance is significant for the colonization and metabolic activity of probiotics to stimulate a positive effect on the host [39]. The survival patterns were determined as 7.40 and 8.18 log CFU/mL for gastric and intestinal fluids, respectively (Fig 7C and D). The resistance of the mentioned stress factors is associated with the existence of the putative genes related to stress resistance, DNA and protein protection/repair and these are listed in Table S15. Especially, cation-proton antiporters (*NapA, NhaC, NhaP*), ABC ATPases, F_0_F_1_-type ATP synthase, and chaperones (Clp ATPases) are responsible for homeostasis and intracellular pH regulation which have contributed to acid and bile resistance. Similarly, the same acid bile resistance mechanisms have been reported in *L. plantarum* Y44 [81], *L. amyloticus* L6 [82], and *L*.*fermentum* YLF016 [39]. On the other side, choloylglycine hydrolase (*cbh*) and inorganic pyrophosphatases (e.g. *PpaC*) were supposed to be prominent in acid bile resistance. Differently, based on antimicrobial activity assay results, CFS of the AGA52 showed a zone of inhibition (> 5 mm) versus all tested pathogen strains except for *S*.*aureus* (Table S16). As far as we are concerned, inhibition of test pathogens by CFS of the AGA52 has originated from the produced organic acids by itself. Because this strain does not encode any bacteriocin production-responsible gene cluster in its genome.

**Fig 7.**
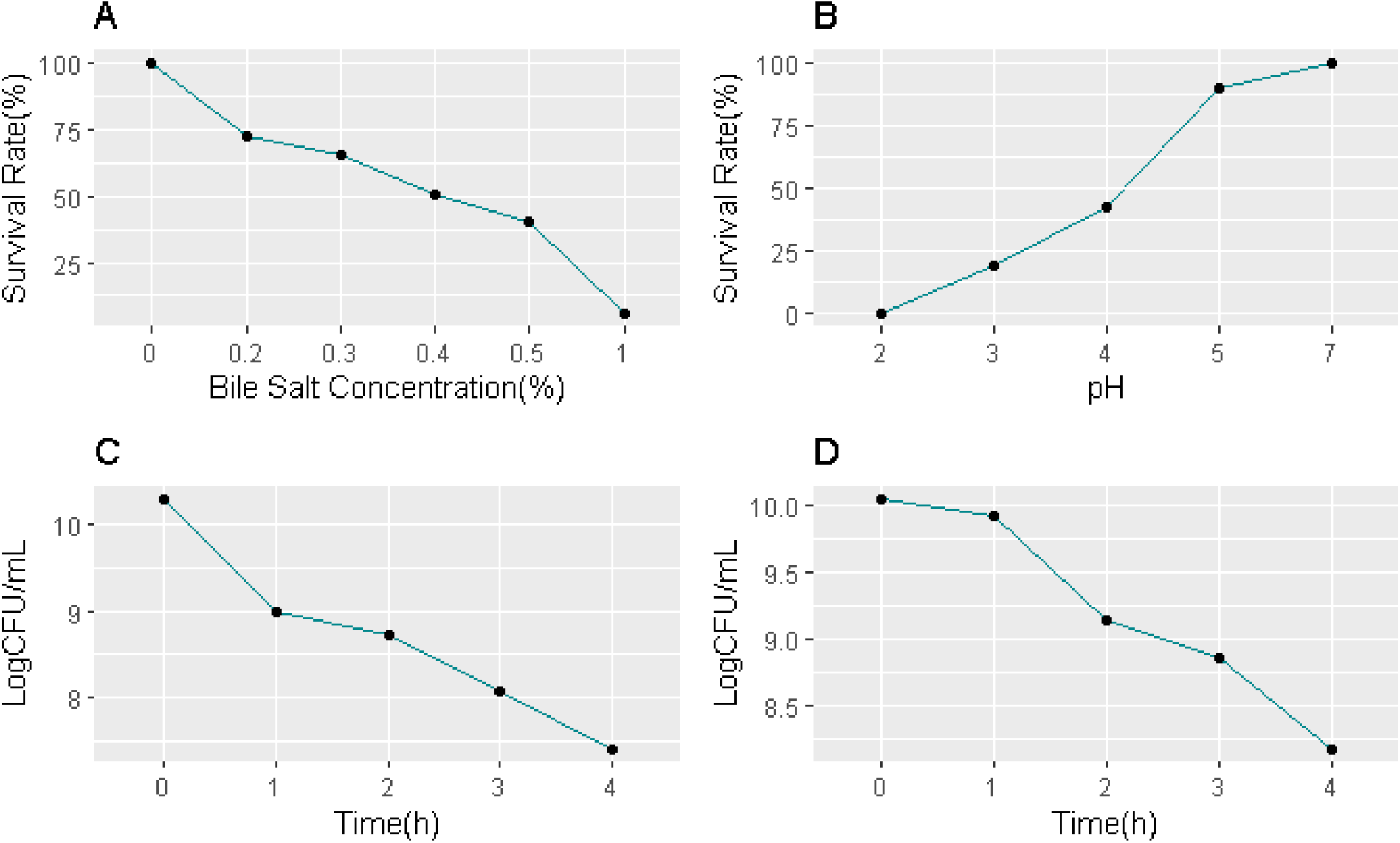
Survival rate and growth characteristics under different conditions. A. Survival at different bile salts concentrations, B. Growth at different pH values, C. Survival tendency of AGA52 at simulated gastric juice environment, D. Survival behaviour of AGA52 at simulated intestinal juice environment.

Antioxidant activity is one of the most important probiotic functions that are directly related to health benefits. Following antioxidant assays, DPPH and ABTS inhibitions were determined as 92.9±0.01% and 85.3±0.065%, respectively. DPPH and ABTS are well-known free radicals which are generally used for testing the free radical scavenging activity of antioxidants and can be analyzed colourimetrically. These strong inhibition values are related to the antioxidant enzymes such as *trxA* (Thioredoxin), *trxB* (Thioredoxin reductase), *tpx* (Thiol peroxidase), *dsbA* (Thiol disulfide oxidoreductase), *arsC* (Arsenate reductase), *Gor* (Glutathione reductase), *nrdH* (Glutaredoxin) and other putative antioxidant genes. The identifiers and putative functions of aforecited genes were summarized in Table S15. The “*in vitro*” and “*in silico*” findings regarding the antioxidant activity of AGA52 are compatible and further demonstrate that it can adapt to higher oxidative stress environments.

### Cholesterol assimilation

Cardiovascular and coronary artery disease risks are directly correlated with dyslipidemia and are important public health concerns. Dyslipidemia generally occurs due to unbalanced concentrations of cholesterol, low-density lipoprotein cholesterol, (LDL-C), triglycerides, and high-density lipoprotein (HDL). Different researchers have reported that various *Lb. fermentum* strains showed blood cholesterol-lowering effects [10, 11]. In this regard, similar cholesterol-decreasing effects have been observed “*in vitro*” for this study. Based on cholesterol assimilation assay results, the AGA52 was assimilated 28.8±0.003% of the 100ppm cholesterol for 24h, while it has been utilized the 33.9±0.005% of total cholesterol added to MRS broth for 48h. In previous studies, 11.51±1.44%, 37.19±4.99%, and 40.84±1.90% cholesterol assimilations were reported for *Lb. fermentum* strains NCIMB 5221, NCIMB 8829, and NCIMB 2797, respectively [83]. In another study, cholesterol assimilation ranges from 28.3-88% were described for different *Lb. fermentum* strains by Pan et al. [84]. The molecular mechanism of hypocholesterolemic effects is generally explained by the presence of bile salt hydrolase (*bsh* or *cbh*) enzymes, or the production of short-chained-fatty acids that inhibit enzymes like 3-hydroxy-3-methylglutaryl coenzyme A which is component of ketogenesis pathways. In this study, the presence of choloylglycine hydrolase (*cbh*) enzyme which has bile salt hydrolase like activity was confirmed “*in silico*”. The presence of organic acids, like propionic, acetic, butyric, and lactic acids was also confirmed by GC-MS (method and data not shown) which is another factor related with cholesterol assimilation.

### GABA production as a psychobiotic feature

Gamma-aminobutyric acid (GABA) is a free amino acid which is produced from L-glutamic acid or MSG by Glutamate decarboxylase (GAD, EC 4.1.1.15) and is extensively found in animals, plants and microbes [85]. GABA has multiple physiological functions and acts as major inhibitory neurotransmitter which delivers chemical message in the central nervous systems of mammalians and has a significant role on stress response, cognition and behaviour [86]. Besides, GABA supplementation is positively affects depression, sleeplessness, immunity, blood pressure regulation, visual cortical function, anxiety, and menopausal syndrome [87-91]. Various lactobacilli strains such as *Lb. fermentum, Lb. plantarum, Lb. helveticus, Lb. paracasei* and *Lb. delbrueckii* sp. *bulgaricus* are able to produce GABA that originated from cheese, pickles, yoghurt, fermented soybean etc. Recently, a great number of Lactic acid bacteria strains have been isolated from various fermented foods and utilized for the enrichment of functionality of foods by supplementation of GABA [85]. In microbes, biosynthesis of GABA is carried out by pyridoxal-5’-phosphate-dependent-glutamic acid decarboxylase (GAD) system which is comprised of the GAD enzyme (encoded by *gadA* or *gadB*) and glutamate/GABA antiporter (*gadC*) [92-94]. In this study, the existence of these genes and GABA production ability from MSG were confirmed via “*in silico*” and “*in vitro*” analyses and the members of GAD system were summarized in Table S15. The NMR spectra were shown in FigS4 describes that the AGA52 can convert the MSG to GABA by using its related enzyme system. It is clearly seen in FigS4 that G1 (CH_2_: 1.76 ppm), G2 (CH_2_: 2.16 ppm), and G3 (CH_2_: 2.87 ppm) peaks of MRS broth are showing highest similartiy with the G1(CH_2_: 1.75 ppm), G2(CH_2_:2.15 ppm), and G3 (CH_2_:2.85 ppm) peaks of GABA standard which is the evidence of the AGA52 is able to produce GABA from MSG.

## Conclusion

Briefly, it has been determined that *Lb. fermentum* AGA52 is a novel strain that possesses crucial probiotic and psychobiotic features such as higher antioxidant activiy, cholesterol assimilation capacity, GABA production ability, auto-aggregation ability, cell surface hydrophobicity, gut persistance, adaptation to gastointestinal simulated environment, acid-bile tolerance. It has obligate heterofermentative carbohydrate metabolism due to the absence of the phosphofructokinase enzyme which is confirmed from its genome. The AGA52 can also produce CO_2_, lactate, and ethanol by heterofermentative pathway. Three phage regions (2 intact and 1 questionable) were identified and were detected as temperate due to the absence of the endolysins and 0.79% of the genome of the AGA52 has orginated from phage related horizontal gene transfer. On the other side, all detected resistome were found as intrinsically originated. The detected CRISPR structures revealed that the AGA52 can be used as model organism for CRISPR studies or can be used for development of next-generation probiotics by acquiring novel functional gene clusters. Besides, the survival behaviour of the strain proved that it has an adaptable nature to gastrointestinal mimicked environment and the diversity of the CAZYmes has also confirmed this case. When desired, survivability of this strain to gastrointestinal conditions can be enhanced more by encapsulation or gaining high copy number of relavant genes (e.g. *bsh*, Na+/H+ antiporters, F_0_F_1_ ATPases). The AGA52 has displayed the highest antioxidant capacity according to DPPH and ABTS+ assays which is also validated by presence of relevant genes. Overall, all findings revealed that the AGA52 has promising bacterial strain which has a potential for use as dietary supplement that might provide functional and therapeutic benefits to the host.

## Acknowledgements

Mr. Mehmet Horzum would like to thank the Council of Higher Education of Türkiye (YÖK) due to the 100/2000 PhD scholarship program.

## Declarations

### Author Contribution

All authors contributed to the study’s conception and design. Material preparation, data collection and analysis were performed by Ahmet E. Yetiman and Mehmet Horzum. The first draft of the manuscript was written by Ahmet E. Yetiman and was proofread by Mikail Akbulut. All authors read and approved the final manuscript.

