## Supplementary materials for "Evaluation of metabolic and functional properties of cholesterol-reducing and GABA-producer *Limosilactobacillus fermentum* strain AGA52 isolated from lactic acid fermented Shalgam by using *in vitro* and *in silico* probiogenomic approaches"

**Table S1.** The Predicted CRISPR systems in the genome of *Limosilactobacillus fermentum* AGA52 (GenBank: CP091132.1)

| Feature Type | Start/End | Strand | Product |
|---|---|---|---|
| Cas9 | 629982/634118 | - | CRISPR-associated endonuclease Cas9 |
| Cas1 | 628869/629774 | - | type II CRISPR-associated endonuclease Cas1 |
| UJP16263.1 | 627918/628589 | - | type II-A CRISPR-associated protein Csn2 |
| UJP16264.1 | 628586/628891 | - | CRISPR-associated endonuclease Cas2 |
| CRISPR-array | 626931/627889 | + | CRISPR region with repeat<br>GTACTCGGAACTACTGATCTGACACTCATCCAAGAC<br>aactctaaaactactgatctgacactcatccaagacaagcatctagatttggttgcttga<br>tttgtggtactcggaactactgatctgacactcatccaagacctaaaggaggaaccgacc<br>atcaatgaactagtactcggaactactgatctgacactcatccaagaccacgacgactga<br>agaactgtacaacaactagtactcggaactactgatctgacactcatccaagactaaagg<br>tttgccgaccaacctaacggacaagtactcggaactactgatctgacactcatccaagac<br>taaaagctatactcaccgtctagatcaagcgtactcggaactactgatctgacactcatc<br>caagacctgatgaacagattgccaaaaatgtccagcgtactcggaactactgatctgaca<br>ctcatccaagacagcaatcgacgagcttaagcgatgatcaaagtactcggaactactgat<br>ctgacactcatccaagacgcaattacgacctttgttgacaaccgcgatgtactcggaact<br>actgatctgacactcatccaagaccggcattgaggactttgtgagctatcacgagtactc<br>ggaactactgatctgacactcatccaagactaatttatcaaagtttgccaacaatgtagg<br>tactcggaactactgatctgacactcatccaagacctcagcccaagtgggcctctacggg<br>agcaagtactcggaactactgatctgacactcatccaagacgactatgcgatgtggatgt<br>atggcgtaaatgtactcggaactactgatctgacactcatccaagacagtcttaccgtcc<br>aatcagagataccgagtgtactcggaactactgatctgacactcatccaagacagaagcc<br>taagtcgccagtttgcacctcacgtactcggaactactgatctgacactcatccaagac |
| CRISPR-spacer | 626967/626996 | + | AAGCATCTAGATTTGGTTGCTTGATTTGTG |
| CRISPR-spacer | 627033/627062 | + | CTAAAGGAGGAACCGACCATCAATGAACTA |
| CRISPR-spacer | 627099/627128 | + | CACGACGACTGAAGAACTGTACAACAACTA |
| CRISPR-spacer | 627165/627194 | + | TAAAGGTTTGCCGACCAACCTAACGGACAA |
| CRISPR-spacer | 627231/627260 | + | TAAAAGCTATACTCACCGTCTAGATCAAGC |
| CRISPR-spacer | 627297/627326 | + | CTGATGAACAGATTGCCAAAAATGTCCAGC |
| CRISPR-spacer | 627363/627392 | + | AGCAATCGACGAGCTTAAGCGATGATCAAA |
| CRISPR-spacer | 627429//627458 | + | GCAATTACGACCTTTGTTGACAACCGCGAT |
| CRISPR-spacer | 627495/627524 | + | CGGCATTGAGGACTTTGTGAGCTATCACGA |
| CRISPR-spacer | 627561/627589 | + | TAATTTATCAAAGTTTGCCAACAATGTAG |
| CRISPR-spacer | 627626/627655 | + | CTCAGCCCAAGTGGGCCTCTACGGGAGCAA |
| CRISPR-spacer | 627692/627721 | + | GACTATGCGATGTGGATGTATGGCGTAAAT |
| CRISPR-spacer | 627758/627758 | + | AGTCTTACCGTCCAATCAGAGATACCGAGT |
| CRISPR-spacer | 627824/627853 | + | AGAAGCCTAAGTCGCCAGTTTGCACCTCAC |
| CRISPR-repeat | 626931/626966 | + | AACTCTAAAACTACTGATCTGACACTCATCCAAGAC |
| CRISPR-repeat | 627854/627889 | + | GTACTCGGAACTACTGATCTGACACTCATCCAAGAC |
| CRISPR-repeat | 627788/627823 | + | GTACTCGGAACTACTGATCTGACACTCATCCAAGAC |
| CRISPR-repeat | 627722/627757 | + | GTACTCGGAACTACTGATCTGACACTCATCCAAGAC |
| CRISPR-repeat | 627656/627691 | + | GTACTCGGAACTACTGATCTGACACTCATCCAAGAC |
| CRISPR-repeat | 627590/627625 | + | GTACTCGGAACTACTGATCTGACACTCATCCAAGAC |
| CRISPR-repeat | 627525/627560 | + | GTACTCGGAACTACTGATCTGACACTCATCCAAGAC |
| CRISPR-repeat | 627459/627494 | + | GTACTCGGAACTACTGATCTGACACTCATCCAAGAC |
| CRISPR-repeat | 627393/627428 | + | GTACTCGGAACTACTGATCTGACACTCATCCAAGAC |
| CRISPR-repeat | 627327/627362 | + | GTACTCGGAACTACTGATCTGACACTCATCCAAGAC |
| CRISPR-repeat | 627261/627296 | + | GTACTCGGAACTACTGATCTGACACTCATCCAAGAC |
| CRISPR-repeat | 627195/627230 | + | GTACTCGGAACTACTGATCTGACACTCATCCAAGAC |
| CRISPR-repeat | 627129/627164 | + | GTACTCGGAACTACTGATCTGACACTCATCCAAGAC |
| CRISPR-repeat | 627063/627098 | + | GTACTCGGAACTACTGATCTGACACTCATCCAAGAC |
| CRISPR-repeat | 626997/627032 | + | GTACTCGGAACTACTGATCTGACACTCATCCAAGAC |

**Table S2.** The predicted prophage regions of *Limosilactobacillus fermentum* strain AGA52

| Region | Length | Completeness | Score | Total Proteins | Region Position | Most Common Phage (Number of matching proteins) | GC% |
|---|---|---|---|---|---|---|---|
| 1 | 13.3 kb | **questionable** | 70 | 15 | 842607-855944 | PHAGE_Klebsi_phiKO2_NC_005857(2) | 42.52% |
| 2 | 32.2 kb | **intact** | 140 | 40 | 918171-951398 | PHAGE_Lactob_LfeSau_NC_029068(16) | 43.70% |
| 3 | 11.7 kb | **intact** | 130 | 16 | 1975601-1987361 | PHAGE_Lactob_521B_NC_048752(2) | 43.94% |

**Table S3.** The questionable and first prophage region elements of *Limosilactobacillus fermentum* AGA52 (PHAGE_Klebsi_phiKO2_NC_005857).

| # | Locus | ORF Start | ORF Stop | Strand | Homolog/Ortholog Species | Homolog/Ortholog Protein | E-Value |
|---|---|---|---|---|---|---|---|
| 1 | PP_00833 | 842607 | 844184 | Backward | Tail protein | PP_00833, tail length tape-measure protein, phage(gi100057), PHAGE_Anoxyb_A403_NC_048701 | 1.13e-27 |
| 2 | PP_00834 | 844177 | 845838 | Backward | Phage-like protein | PP_00834, site-specific serine recombinase, phage(gi89152504), PHAGE_Bacill_Fah_NC_007814 | 9.71e-35 |
| 3 | PP_00835 | 845874 | 846107 | Backward | Hypothetical protein | PP_00835, hypothetical | N/A |
| 4 | PP_00836 | 846200 | 847078 | Backward | Hypothetical protein | PP_00836, hypothetical protein, phage(gi100039), PHAGE_Faecal_FP_Lugh_NC_047912 | 5.01e-30 |
| 5 | PP_00837 | 847075 | 847488 | Backward | Phage-like protein | PP_00837, holin, phage(gi89152494), PHAGE_Bacill_Fah_NC_007814 | 4.46e-29 |
| 6 | PP_00838 | 847475 | 847858 | Backward | Head protein | PP_00838, head-tail adaptor protein, phage(gi971821549), PHAGE_Clostr_phiCT19406C_NC_029006 | 1.41e-13 |
| 7 | PP_00839 | 848153 | 849331 | Backward | Head protein | PP_00839, minor capsid protein, phage(gi589893767), PHAGE_Geobac_GBK2_NC_023612 | 2.25e-44 |
| 8 | PP_00840 | 849351 | 850022 | Backward | Phage-like protein | PP_00840, putative Clp peptidase, phage(gi148747731), PHAGE_Geobac_E2_NC_009552 | 1.65e-46 |
| 9 | PP_00841 | 850019 | 851284 | Backward | Portal protein | PP_00841, putative portal protein, phage(gi46402090), PHAGE_Klebsi_phiKO2_NC_005857 | 1.17e-86 |
| 10 | PP_00842 | 851313 | 852911 | Backward | Hypothetical protein | PP_00842, hypothetical protein, phage(gi100002), PHAGE_Gordon_Nymphadora_NC_031061 | 2.96e-105 |
| 11 | PP_00843 | 852975 | 853181 | Backward | Hypothetical protein | PP_00843, hypothetical | N/A |
| 12 | PP_00844 | 853184 | 853804 | Backward | Hypothetical protein | PP_00844, hypothetical | N/A |
| 13 | PP_00845 | 853877 | 855103 | Backward | Head protein | PP_00845, head morphogenesis protein, phage(gi100068), PHAGE_Bacill_vB_BtS_B83_NC_048762 | 4.29e-140 |
| 14 | PP_00846 | 855103 | 855645 | Backward | Hypothetical protein | PP_00846, hypothetical | N/A |
| 15 | PP_00847 | 855765 | 855944 | Backward | Phage-like protein | PP_00847, Gp60, phage(gi46402146), PHAGE_Klebsi_phiKO2_NC_005857 | 1.61e-07 |

**Table S4.** The second prophage (intact) region elements of *Limosilactobacillus fermentum* AGA52 (PHAGE_Lactob_LfeSau_NC_029068).

| # | Locus | ORF Start | ORF Stop | Strand | Homolog/Ortholog Species | Homolog/Ortholog Protein | E-Value |
|---|---|---|---|---|---|---|---|
| 1 | PP_00908 | 918171 | 923060 | Backward | Phage-like protein | PP_00908, structural protein, phage(gi985757742), PHAGE_Lactob_LfeSau_NC_029068 | 0.0 |
| 2 | attL | 922221 | 922233 | Forward | Attachment site (AAAAACAGTTAAG) | attL | N/A |
| 3 | PP_00909 | 923057 | 923842 | Backward | Tail protein | PP_00909, tail protein, phage(gi985757741), PHAGE_Lactob_LfeSau_NC_029068 | 5.17e-82 |
| 4 | PP_00910 | 923855 | 926965 | Backward | Phage-like protein | PP_00910, tape measure protein, phage(gi985757740), PHAGE_Lactob_LfeSau_NC_029068 | 0.0 |
| 5 | PP_00911 | 927395 | 927727 | Backward | Phage-like protein | PP_00911, putative tapemeasure chaperone protein, phage(gi985757738), PHAGE_Lactob_LfeSau_NC_029068 | 1.41e-54 |
| 6 | PP_00912 | 927742 | 928332 | Backward | Tail protein | PP_00912, major tail protein, phage(gi985757737), PHAGE_Lactob_LfeSau_NC_029068 | 6.09e-113 |
| 7 | PP_00913 | 928348 | 928746 | Backward | Hypothetical protein | PP_00913, hypothetical protein, phage(gi985757736), PHAGE_Lactob_LfeSau_NC_029068 | 1.79e-59 |
| 8 | PP_00914 | 928743 | 928889 | Backward | Head protein | PP_00914, head/tail component, phage(gi985757735), PHAGE_Lactob_LfeSau_NC_029068 | 2.78e-26 |
| 9 | PP_00915 | 929390 | 929776 | Backward | Phage-like protein | PP_00915, putative DNA packaging protein, phage(gi985757733), PHAGE_Lactob_LfeSau_NC_029068 | 7.57e-53 |
| 10 | PP_00916 | 929933 | 930796 | Backward | Head protein | PP_00916, capsid protein, phage(gi985757732), PHAGE_Lactob_LfeSau_NC_029068 | 3.19e-157 |
| 11 | PP_00917 | 930809 | 931411 | Backward | Hypothetical protein | PP_00917, hypothetical protein, phage(gi985757731), PHAGE_Lactob_LfeSau_NC_029068 | 1.09e-92 |
| 12 | PP_00918 | 931772 | 932773 | Backward | Head protein | PP_00918, head morphogenesis protein, phage(gi985757729), PHAGE_Lactob_LfeSau_NC_029068 | 1.10e-178 |
| 13 | PP_00919 | 932754 | 934139 | Backward | Portal protein | PP_00919, portal protein, phage(gi985757728), PHAGE_Lactob_LfeSau_NC_029068 | 0.0 |
| 14 | PP_00920 | 934152 | 935231 | Backward | Terminase | PP_00920, terminase large subunit, phage(gi985757727), PHAGE_Lactob_LfeSau_NC_029068 | 0.0 |
| 15 | PP_00921 | 935546 | 935953 | Backward | Terminase | PP_00921, putative terminase small subunit, phage(gi571797877), PHAGE_Lactob_phiJB_NC_022775 | 2.68e-32 |
| 16 | PP_00922 | 936109 | 936699 | Backward | Hypothetical protein | PP_00922, hypothetical protein, phage(gi281416358), PHAGE_Entero_phiFL1A_NC_013646 | 5.59e-31 |
| 17 | PP_00923 | 936884 | 937402 | Backward | Phage-like protein | PP_00923, transcriptional regulator, phage(gi985757774), PHAGE_Lactob_LfeSau_NC_029068 | 1.31e-08 |
| 18 | PP_00924 | 937580 | 938032 | Backward | Hypothetical protein | PP_00924, hypothetical | N/A |
| 19 | PP_00925 | 938061 | 938264 | Backward | Hypothetical protein | PP_00925, hypothetical | N/A |
| 20 | PP_00926 | 938288 | 938770 | Backward | Tail protein | PP_00926, tail length tape-measure protein, phage(gi100057), PHAGE_Lactob_Lenus_NC_047897 | 1.12e-29 |
| 21 | PP_00927 | 939435 | 939617 | Backward | Hypothetical protein | PP_00927, hypothetical protein, phage(gi937456199), PHAGE_Lactob_phiPYB5_NC_027982 | 1.10e-17 |
| 22 | PP_00928 | 939621 | 940412 | Backward | Phage-like protein | PP_00928, DNA replication protein, phage(gi418489432), PHAGE_Lactob_LF1_NC_019486 | 8.80e-123 |
| 23 | PP_00929 | 940421 | 940729 | Backward | Phage-like protein | PP_00929, putative phage replication protein, phage(gi937456197), PHAGE_Lactob_phiPYB5_NC_027982 | 6.32e-49 |
| 24 | PP_00930 | 941431 | 941646 | Backward | Hypothetical protein | PP_00930, hypothetical | N/A |
| 25 | PP_00931 | 941660 | 942145 | Backward | Phage-like protein | PP_00931, single-strand binding protein, phage(gi418489430), PHAGE_Lactob_LF1_NC_019486 | 1.12e-73 |
| 26 | PP_00932 | 942146 | 942994 | Backward | Hypothetical protein | PP_00932, hypothetical protein, phage(gi418489429), PHAGE_Lactob_LF1_NC_019486 | 0.0 |
| 27 | PP_00933 | 942987 | 943982 | Backward | Phage-like protein | PP_00933, RecT protein, phage(gi418489428), PHAGE_Lactob_LF1_NC_019486 | 1.76e-157 |
| 28 | PP_00934 | 943985 | 944194 | Backward | Hypothetical protein | PP_00934, hypothetical protein, phage(gi985757758), PHAGE_Lactob_LfeSau_NC_029068 | 1.11e-13 |
| 29 | PP_00935 | 944229 | 944390 | Backward | Hypothetical protein | PP_00935, hypothetical | N/A |

**Table S4.** The second prophage (intact) region elements of *Limosilactobacillus fermentum* AGA52 (continues)

| # | Locus | ORF Start | ORF Stop | Strand | Homolog/Ortholog Species | Homolog/Ortholog Protein | E-Value |
|---|---|---|---|---|---|---|---|
| 30 | PP_00936 | 944405 | 944716 | Backward | Hypothetical protein | PP_00936, hypothetical protein, phage(gi148750868), PHAGE_Lactob_LL_H_NC_009554 | 6.02e-08 |
| 31 | PP_00937 | 944831 | 945088 | Backward | Hypothetical protein | PP_00937, hypothetical protein, phage(gi971767221), PHAGE_Brevib_Abouo_NC_029029 | 1.27e-07 |
| 32 | PP_00938 | 945112 | 945300 | Backward | Phage-like protein | PP_00938, repressor, phage(gi13095748), PHAGE_Lactoc_bIL286_NC_002667 | 7.61e-14 |
| 33 | PP_00939 | 945502 | 945837 | Forward | Phage-like protein | PP_00939, cl-like repressor phage associated, phage(gi588295084), PHAGE_Strept_20617_NC_023503 | 1.37e-26 |
| 34 | PP_00940 | 945842 | 946240 | Forward | Phage-like protein | PP_00940, Lj965 prophage repressor-like protein, phage(gi418489418), PHAGE_Lactob_LF1_NC_019486 | 8.06e-39 |
| 35 | PP_00941 | 946426 | 946737 | Forward | Hypothetical protein | PP_00941, hypothetical | N/A |
| 36 | PP_00942 | 946835 | 947326 | Forward | Hypothetical protein | PP_00942, hypothetical | N/A |
| 37 | attR | 947491 | 947503 | Forward | Attachment site (AAAAACAGTTAAG) | attR | N/A |
| 38 | PP_00943 | 947549 | 947692 | Forward | Hypothetical protein | PP_00943, hypothetical protein, phage(gi41179289), PHAGE_Lactob_Lj928_NC_005354 | 1.42e-15 |
| **39** | **PP_00944** | **947854** | **948942** | **Forward** | **Integrase** | **PP_00944, phage integrase, phage(gi418489411), PHAGE_Lactob_LF1_NC_019486** | **4.16e-106** |
| 40 | PP_00945 | 949203 | 949823 | Backward | Hypothetical protein | PP_00945, hypothetical | N/A |
| 41 | PP_00946 | 949941 | 950387 | Forward | Terminase | PP_00946, terminase small subunit, phage(gi100071), PHAGE_Lactob_8014_B2_NC_047739 | 1.64e-21 |
| 42 | PP_00947 | 950409 | 951398 | Forward | Hypothetical protein | PP_00947, hypothetical protein, phage(gi100002), PHAGE_Entero_DE3_NC_042057 | 1.96e-14 |

**Table S5.** The third prophage (intact) region elements of *Limosilactobacillus fermentum* AGA52 (PHAGE_Lactob_521B_NC_048752).

| # | Locus | ORF Start | ORF Stop | Strand | Homolog/Ortholog Species | Homolog/Ortholog Protein | E-Value |
|---|---|---|---|---|---|---|---|
| 1 | PP_01962 | 1975601 | 1976590 | Backward | Transposase | **PP_01962**, transposase, phage(gi56693108), PHAGE_Lactob_LP65_NC_006565 | 1.85e-156 |
| 2 | PP_01963 | 1976717 | 1977595 | Backward | Transposase | **PP_01963**, transposase, phage(gi971752662), PHAGE_Lactob_CL1_NC_028888 | 5.24e-106 |
| 3 | PP_01964 | 1977595 | 1977735 | Backward | Head protein | PP_01964, head morphogenesis protein, phage(gi100068), PHAGE_Bacill_Shbh1_NC_030925 | 6.73e-16 |
| 4 | PP_01965 | 1977834 | 1978859 | Forward | Transposase | **PP_01965**, IS30 family transposase, phage(gi15677611), PROPHAGE_Neisse_MC58 | 2.59e-38 |
| 5 | PP_01966 | 1978937 | 1979911 | Forward | Transposase | **PP_01966**, putative transposase, phage(gi588498272), PHAGE_Staphy_StauST398_4_NC_023499 | 8.43e-62 |
| 6 | PP_01967 | 1980735 | 1980872 | Backward | Hypothetical protein | PP_01967, hypothetical | N/A |
| 7 | PP_01968 | 1981118 | 1981669 | Backward | Transposase | **PP_01968**, transposase, phage(gi971743568), PHAGE_Bacter_Diva_NC_028788 | 9.61e-25 |
| 8 | PP_01970 | 1981762 | 1981914 | Backward | Transposase | **PP_01970**, transposase, phage(gi971756981), PHAGE_Paenib_Tripp_NC_028930 | 1.57e-08 |
| 9 | PP_01969 | 1982077 | 1982211 | Forward | Hypothetical protein | PP_01969, hypothetical | N/A |
| 10 | PP_01971 | 1982637 | 1983335 | Forward | Hypothetical protein | PP_01971, hypothetical | N/A |
| 11 | PP_01972 | 1983316 | 1983447 | Forward | Transposase | **PP_01972**, transposase, phage(gi971741608), PHAGE_Paenib_Vegas_NC_028767 | 2.52e-06 |
| 12 | PP_01973 | 1983673 | 1984347 | Forward | Transposase | **PP_01973**, putative transposase, phage(gi9626114), PHAGE_Spirop_1_R8A2B_NC_001365 | 1.11e-11 |
| 13 | PP_01974 | 1984362 | 1984577 | Forward | Hypothetical protein | PP_01974, hypothetical protein, phage(gi371496264), PHAGE_Plankt_PaV_LD_NC_016564 | 5.48e-05 |
| 14 | PP_01975 | 1984601 | 1985197 | Forward | Transposase | **PP_01975**, ISxac3 transposase, phage(gi21231086), PROPHAGE_Xantho_33913 | 6.59e-22 |
| 15 | PP_01976 | 1985992 | 1986651 | Backward | Transposase | **PP_01976**, transposase, phage(gi971752662), PHAGE_Lactob_CL1_NC_028888 | 1.35e-47 |
| 16 | PP_01977 | 1986642 | 1987361 | Backward | Transposase | **PP_01977**, putative transposase, phage(gi588498272), PHAGE_Staphy_StauST398_4_NC_023499 | 5.41e-19 |

**Table S6.** Horizontal gene transfer confirmation of the strain AGA52 according to ProteinBLAST results of prophage regions

| S/N | # | Locus | ORF Start | ORF Stop | Strand | Homolog/Ortholog Species | Protein BLAST | Accession | E-value |
|---|---|---|---|---|---|---|---|---|---|
| **The questionable and first prophage region elements of *Limosilactobacillus fermentum* strain AGA52** | | | | | | | | | |
| 1 | 1 | PP_00833 | 842607 | 844184 | Backward | Tail protein | recombinase family protein [***Ligilactobacillus animalis***] | **WP_216547624.1** | 0.0 |
| 2 | 2 | PP_00834 | 844177 | 845838 | Backward | Phage-like protein | MULTISPECIES: recombinase family protein [***Lactobacillaceae]*** | WP_094537226.1 | 0.0 |
| 3 | 4 | PP_00836 | 846200 | 847078 | Backward | Hypothetical protein | 1,4-beta-N-acetylmuramidase [***Pediococcus acidilactici***] | **WP_159216384.1** | 0.0 |
| 4 | 5 | PP_00837 | 847075 | 847488 | Backward | Phage-like protein | phage holin family protein [***Limosilactobacillus reuteri***] | **WP_094537229.1** | 2,00E-91 |
| 5 | 7 | PP_00839 | 848153 | 849331 | Backward | Head protein | phage major capsid protein [***Pediococcus acidilactici***] | **WP_159216387.1** | 0.0 |
| 6 | 9 | PP_00841 | 850019 | 851284 | Backward | Portal protein | phage portal protein [***Pediococcus acidilactici***] | **WP_200835729.1** | 0.0 |
| 7 | 10 | PP_00842 | 851313 | 852911 | Backward | Hypothetical protein | putative phage terminase, large subunit [***Limosilactobacillus reuteri* SD2112**] | **AEI56698.1** | 0.0 |
| 8 | 11 | PP_00843 | 852975 | 853181 | Backward | Hypothetical protein | MULTISPECIES: DUF5049 domain-containing protein [***Lactobacillaceae***] | WP_003671984.1 | 1,00E-40 |
| 9 | 12 | PP_00844 | 853184 | 853804 | Backward | Hypothetical protein | MULTISPECIES: hypothetical protein [***Lactobacillaceae***] | WP_075140147.1 | 3,00E-151 |
| 10 | 13 | PP_00845 | 853877 | 855103 | Backward | Head protein | MULTISPECIES: site-specific DNA-methyltransferase [***Lactobacillaceae***] | WP_003671982.1 | 0.0 |
| 11 | 14 | PP_00846 | 855103 | 855645 | Backward | Hypothetical protein | MULTISPECIES: P27 family phage terminase small subunit [***Lactobacillaceae*]** | WP_003671981.1 | 9,00E-133 |
| 12 | 15 | PP_00847 | 855765 | 855944 | Backward | Phage-like protein | MULTISPECIES: HNH endonuclease [***Lactobacillaceae]*** | WP_079376308.1 | 6,00E-35 |
| **The second prophage (intact) region elements of *Limosilactobacillus fermentum* strain AGA52** | | | | | | | | | |
| 13 | 1 | PP_00908 | 918171 | 923060 | Backward | Phage-like protein | TPA: MAG TPA: Minor structural protein 4 [***Siphoviridae* sp.**] | **DAT42420.1** | 0.0 |
| 14 | 4 | PP_00910 | 923855 | 926965 | Backward | Phage-like protein | TPA: MAG TPA: minor tail protein [***Siphoviridae* sp.**] | **DAO43398.1** | 0.0 |
| 15 | 8 | PP_00914 | 928743 | 928889 | Backward | Head protein | hypothetical protein [***Limosilactobacillus reuteri***] | **MRH08483.1** | 7E-27 |
| **The third prophage (intact) region elements of *Limosilactobacillus fermentum* strain AGA52** | | | | | | | | | |
| 16 | 7 | PP_01968 | 1981118 | 1981669 | Backward | Transposase | MULTISPECIES: IS5 family transposase [***Lentilactobacillus***] | **WP_225425087.1** | 3E-27 |

**Table S7.** The predicted transposases of the *Limosilactobacillus fermentum* strain AGA52 genome by using IS Finder

| # | Sequences producing significant alignments | IS Family | Group | Origin | Score (bits) | E-value |
|---|---|---|---|---|---|---|
| 1 | ISLpl2 | IS3 | IS150 | *Lactobacillus plantarum* | 2438 | 0.0 |
| 2 | ISL1 | IS3 | IS3 | *Lactobacillus casei* | 2405 | 0.0 |
| 3 | IS1163 | IS3 | IS3 | *Lactobacillus sake* | 2252 | 0.0 |
| 4 | ISLhe30 | IS30 | | *Lactobacillus helveticus* | 1580 | 0.0 |
| 5 | ISEfm1 | IS982 | | *Enterococcus faecium* | 1542 | 0.0 |
| 6 | ISLca1 | IS3 | IS150 | *Lactobacillus casei* | 1439 | 0.0 |
| 7 | IS19 | IS982 | | *Lactococcus lactis* | 1409 | 0.0 |
| 8 | IS153 | IS3 | IS3 | *Lactobacillus sanfranciscensis* | 977 | 0.0 |
| 9 | ISLasa2 | IS3 | IS150 | *Lactobacillus salivarius* | 153 | 1,00E-33 |
| 10 | ISLhe65 | IS200/IS605 | IS1341 | *Lactobacillus helveticus* | 89.7 | 2,00E-14 |
| 11 | ISLpl1 | IS30 | | *Lactobacillus plantarum* | 83.8 | 1,00E-12 |
| 12 | ISXne2 | IS6 | | *Xenorhabdus nematophila* | 75.8 | 3,00E-10 |
| 13 | ISLjo5 | IS200/IS605 | IS605 | *Lactobacillus johnsonii* | 69.9 | 2,00E-08 |
| 14 | ISMmu1 | IS200/IS605 | IS605 | *Mitsuokella multacida* | 63.9 | 1,00E-06 |
| 15 | ISLre1 | IS4 | ISPepr1 | *Lactobacillus reuteri* | 63.9 | 1,00E-06 |
| 16 | IS946V | IS6 | | *Lactococcus lactis* | 61.9 | 4,00E-06 |
| 17 | IS240F | IS6 | | *Bacillus thuringiensis* | 60.0 | 2,00E-05 |
| 18 | ISBth20 | IS6 | | *Bacillus thuringiensis* | 58.0 | 6,00E-05 |
| 19 | ISS1Z | IS6 | | *Lactococcus lactis* | 58.0 | 6,00E-05 |
| 20 | ISS1X | IS6 | | *Lactococcus lactis* | 58.0 | 6,00E-05 |
| 21 | ISS1T | IS6 | | *Lactococcus lactis* | 58.0 | 6,00E-05 |
| 22 | ISS1S | IS6 | | *Lactococcus lactis* | 58.0 | 6,00E-05 |
| 23 | ISS1RS | IS6 | | *Lactococcus lactis* | 58.0 | 6,00E-05 |
| 24 | ISS1B | IS6 | | *Lactococcus lactis* | 58.0 | 6,00E-05 |
| 25 | ISS1A | IS6 | | *Lactococcus lactis* | 58.0 | 6,00E-05 |
| 26 | ISPp1 | IS30 | | *Pediococcus pentosaceus* | 58.0 | 6,00E-05 |
| 27 | ISEcl11 | IS30 | | *Enterobacter cloacae* | 56.0 | 2,00E-04 |
| 28 | ISHar2 | IS3 | IS3 | *Herminiimonas arsenicoxydans* | 54.0 | 0.001 |
| 29 | ISEfa4 | IS200/IS605 | IS605 | *Enterococcus faecium* | 54.0 | 0.001 |
| 30 | ISEfa9 | IS3 | IS3 | *Enterococcus faecium* | 52.0 | 0.004 |
| 31 | ISDet4 | IS256 | | *Dehalococcoides ethenogenes* | 52.0 | 0.004 |
| 32 | ISSne1 | IS256 | | *Sporosarcina newyorkensis* | 50.1 | 0.014 |
| 33 | ISAcl1 | IS30 | | *Arthrobacter chlorophenolicus* | 50.1 | 0.014 |
| 34 | ISRde1 | IS3 | IS51 | *Roseobacter denitrificans* | 50.1 | 0.014 |
| 35 | ISLiv2 | IS256 | | *Listeria ivanovii* | 48.1 | 0.05 |
| 36 | MITEBth2 | IS6 | | *Bacillus thuringiensis* | 48.1 | 0.05 |
| 37 | ISSsu6 | ISL3 | | *Streptococcus suis* | 48.1 | 0.05 |
| 38 | ISLgar5 | IS256 | | *Lactococcus garvieae* | 48.1 | 0.05 |
| 39 | ISEfm2 | IS256 | | *Enterococcus faecium* | 48.1 | 0.05 |
| 40 | ISEfa13 | IS256 | | *Enterococcus faecium* | 48.1 | 0.05 |
| 41 | ISLhe2 | ISL3 | | *Lactobacillus helveticus* | 48.1 | 0.05 |
| 42 | ISEf1 | IS256 | | *Enterococcus faecalis* | 48.1 | 0.05 |
| 43 | ISRme13 | IS3 | IS3 | *Ralstonia metallidurans* | 48.1 | 0.05 |
| 44 | ISAzvi9 | IS3 | IS3 | *Azotobacter vinelandii* | 48.1 | 0.05 |
| 45 | IS6770 | IS30 | | *Enterococcus faecalis* | 48.1 | 0.05 |

**Table S8.** Antibiotic susceptibility test results with resistome search matches from KofamKOALA and PATRIC 3.6.12.

| Antibiotic Group | Antibiotic | Inhibition zone diameter/status | Antibiotic Resistance Genes | | | | | |
|---|---|---|---|---|---|---|---|---|
| | | | KofamKOALA (ver. 2022-03-01, KEGG release 101.0) | | | PATRIC 3.6.12. | | |
| | | | Gene | Product | E-value | Gene | Product | E-value |
| β-Lactams | Ampicillin (10 μg) | 36.41mm **(S)** | - | - | - | - | - | - |
| | Methicillin (5 μg) | **≤14mm (R)** | *blaI* | BlaI family transcriptional regulator, penicillinase repressor | 7e-36 | - | - | - |
| | | | *mrcA* | penicillin-binding protein 1A [EC:2.4.1.129 3.4.16.4] | 1.6e-193 | | | |
| | | | *pbp2A* | penicillin-binding protein 2A [EC:2.4.1.129 3.4.16.4] | 2.4e-283 | | | |
| | Oxacillin (1 μg) | **≤14mm (R)** | *penP* | beta-lactamase class A [EC:3.5.2.6] | 3.2e-29 | | | |
| | Penicillin G (10 U) | 36.03mm **(S)** | - | - | - | - | - | - |
| | Carbenicillin (100 μg) | 41.52mm **(S)** | - | - | - | - | - | - |
| | Amoxycillin (25 μg) | 33.55mm **(S)** | - | - | - | - | - | - |
| Aminoglycosides | Streptomycin (10 μg) | **≤14mm (R)** | - | - | - | *gidB* | 16S rRNA (guanine(527)-N(7))-methyltransferase (EC 2.1.1.170) | 0.0 |
| | Vancomycin ( 30 μg) | **≤14mm (R)** | *mraY* | phospho-N-acetylmuramoyl-pentapeptide-transferase [EC:2.7.8.13] | 3.1e-114 | - | - | - |
| | | | *alr* | alanine racemase [EC:5.1.1.1] | 3.5e-127 | | | |
| | | | *ddl* | D-alanine-D-alanine ligase [EC:6.3.2.4] | 3.7e-108 | | | |
| | | | *murF* | UDP-N-acetylmuramoyl-tripeptide--D-alanyl-D-alanine ligase [EC:6.3.2.10] | 1.5e-143 | | | |
| | | | *murG* | UDP-N-acetylglucosamine--N-acetylmuramyl-(pentapeptide) pyrophosphoryl-undecaprenol N-acetylglucosamine transferase [EC:2.4.1.227] | 6.1e-124 | | | |
| | | | *vanSC, vanSE, vanSG* | two-component system, OmpR family, sensor histidine kinase VanS [EC:2.7.13.3] | 1.3e-123 | | | |
| | Amikacin (30 μg) | ≤14mm **(R)** | - | - | - | - | - | - |
| | Kanamycin (30 μg) | ≤14mm **(R)** | - | - | - | - | - | - |
| Macrolides | Azithromycin (15 μg) | 16.82 **(I)** | - | - | - | - | - | - |
| | Erythromycin (10 μg) | 18.75 **(I)** | - | - | - | - | - | - |
| Tetracyclines | Tetracycline (30 μg) | 19.7mm **(I)** | - | - | - | - | - | - |
| Rifamycins | Rifampicin (5 μg) | 21.54mm **(S)** | - | - | - | - | - | - |
| Peptide antibiotics | Daptomycin | **Not tested** | - | - | - | *pgsA* | CDP-diacylglycerol--glycerol-3-phosphate 3-phosphatidyltransferase (EC 2.7.8.5) | 1e-134 |
| | | | - | - | - | *mprF* | L-O-lysylphosphatidylglycerol synthase (EC 2.3.2.3) | 0.0 |
| | | | - | - | - | gdpD | Glycerophosphoryl diester phosphodiesterase (EC 3.1.4.46) | 0.0 |

**Table S9.** Horizontal gene transfer screening results for predicted antibiotic-resistance genes of *Limosilactobacillus fermentum* strain AGA52

| Antibiotic | Gene | Protein BLAST result | Accession | Identity | E-Value |
|---|---|---|---|---|---|
| β-Lactams | *blaI* | CopY/TcrY family copper transport repressor [***Limosilactobacillus fermentum***] | WP_012391310.1 | 100% | 2e-107 |
| | *blaI* | CopY/TcrY family copper transport repressor [***Limosilactobacillus fermentum***] | BAW87705.1 | 100% | 5e-102 |
| | *mrcA* | transglycosylase domain-containing protein [***Limosilactobacillus fermentum***] | WP_004563220.1 | 100% | 0.0 |
| | *pbp2A* | PBP1A family penicillin-binding protein [***Limosilactobacillus fermentum***] | WP_014562557.1 | 99% | 0.0 |
| | *penP* | serine hydrolase [***Limosilactobacillus fermentum***] | WP_046025459.1 | 100% | 0.0 |
| Streptomycin | *gidB* | 16S rRNA methyltransferase GidB [***Limosilactobacillus fermentum* F-6**] | AGL87999.1 | 100% | 9e-174 |
| Vancomycin | *mraY* | phospho-N-acetylmuramoyl-pentapeptide-transferase [***Limosilactobacillus fermentum***] | WP_070955466.1 | 100% | 0.0 |
| | *alr* | alanine racemase [***Limosilactobacillus fermentum***] | WP_003684030.1 | 100% | 0.0 |
| | *ddl* | D-alanine--D-alanine ligase [***Limosilactobacillus fermentum***] | WP_003685886.1 | 100% | 0.0 |
| | *murF* | UDP-N-acetylmuramoyl-tripeptide--D-alanyl-D-alanine ligase [***Limosilactobacillus fermentum***] | WP_003686468.1 | 100% | 0.0 |
| | *murG* | UDP-N-acetylglucosamine--N-acetylmuramyl-(pentapeptide) pyrophosphoryl-undecaprenol N-acetylglucosamine transferase [***Limosilactobacillus fermentum***] | GIC73609.1 | 100% | 0.0 |
| | *vanSC, vanSE, vanSG* | HAMP domain-containing histidine kinase [***Limosilactobacillus fermentum***] | WP_100184325.1 | 100% | 0.0 |
| Daptomycin | *pgsA* | CDP-diacylglycerol--glycerol-3-phosphate 3-phosphatidyltransferase [***Limosilactobacillus fermentum***] | WP_003686012.1 | 100% | 2e-124 |
| | *mprF* | bifunctional lysylphosphatidylglycerol flippase/synthetase MprF [***Limosilactobacillus fermentum***] | WP_187703766.1 | 100% | 0.0 |
| | *gdpD* | glycerophosphodiester phosphodiesterase [***Limosilactobacillus fermentum***] | WP_111523276.1 | 100% | 2e-166 |
| Efflux pumps | *efrA, efrE* | ABC transporter ATP-binding protein/permease [***Limosilactobacillus fermentum***] | WP_088460398.1 | 99% | 0.0 |
| | *efrB, efrF* | ABC transporter transmembrane domain-containing protein [***Limosilactobacillus fermentum***] | WP_236098087.1 | 100% | 0.0 |

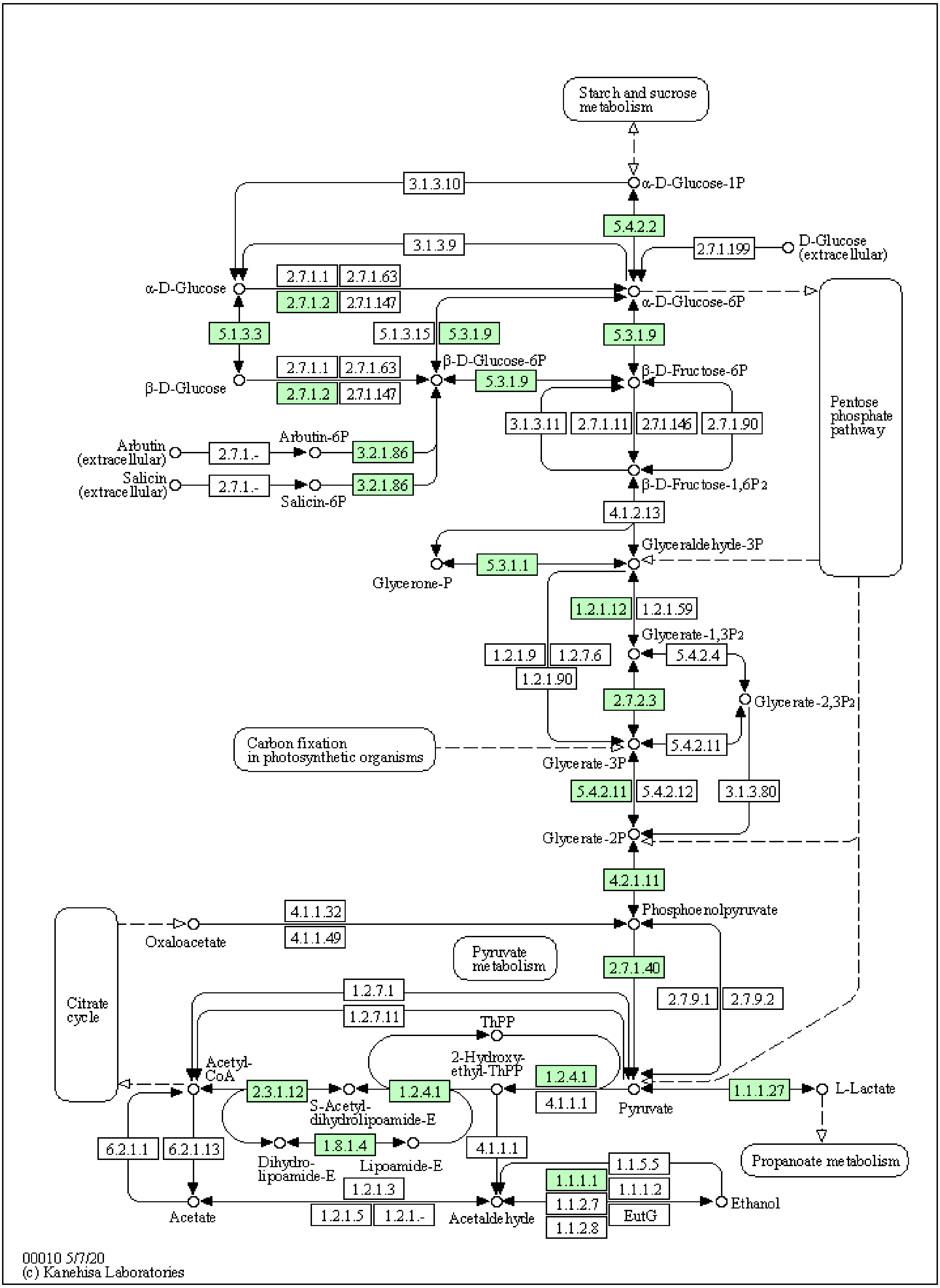


**Fig S1.** The graphical presentation of the possessed enzymes in **glycolysis/gluconeogenesis (EMP)** pathways of limosilactobacillus fermentum AGA52 was obtained from KEGG Mapper (Green coloured EC numbers indicate the presence of the pathway enzymes).

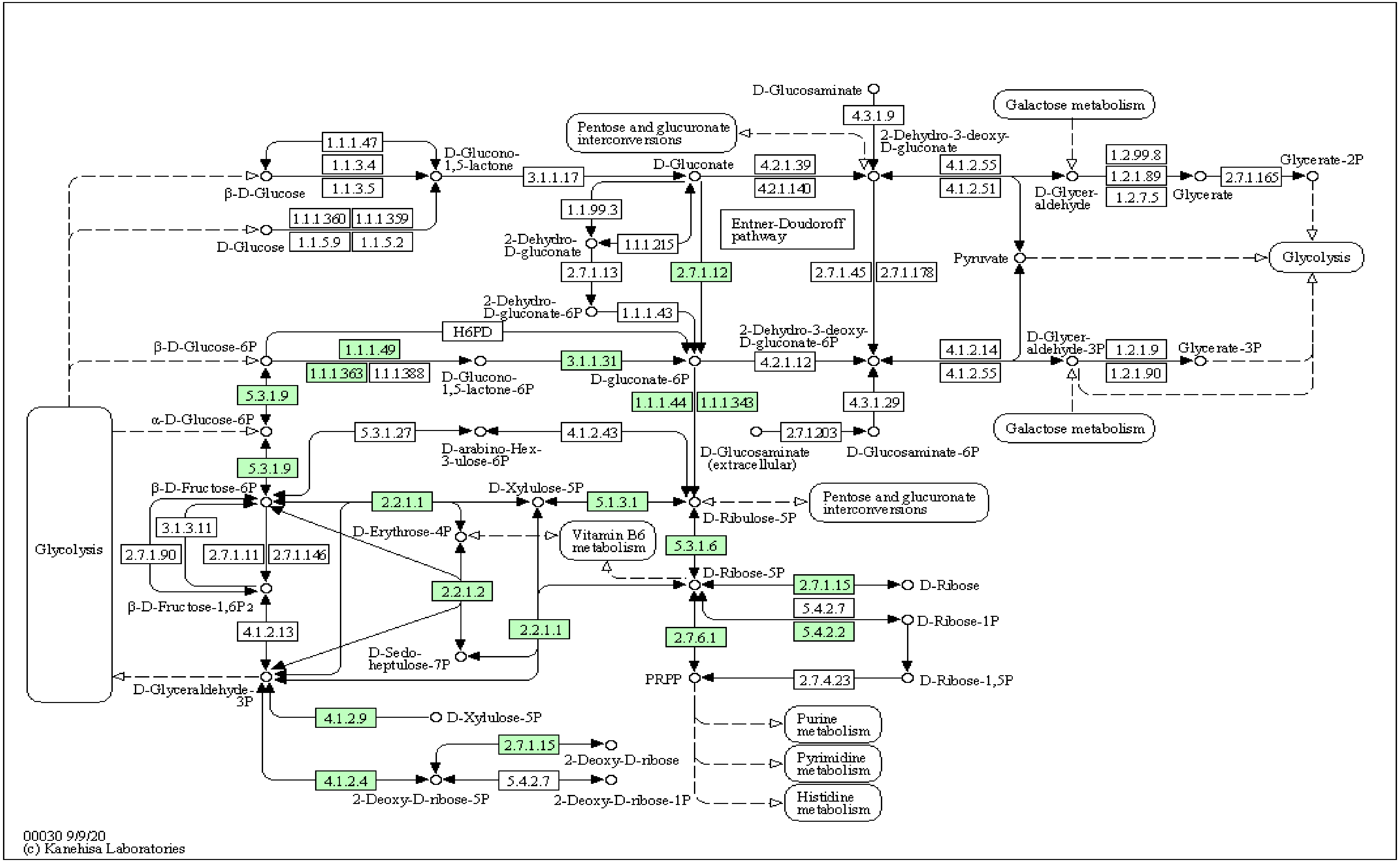


**Fig S2.** The graphical presentation of **pentose phosphate (phospoketolase) pathway** (phospoketolase) enzymes of *Limosilactobacillus fermentum* AGA52 was obtained from KEGG Mapper (Green coloured EC number indicates the presence of the pathway enzymes).

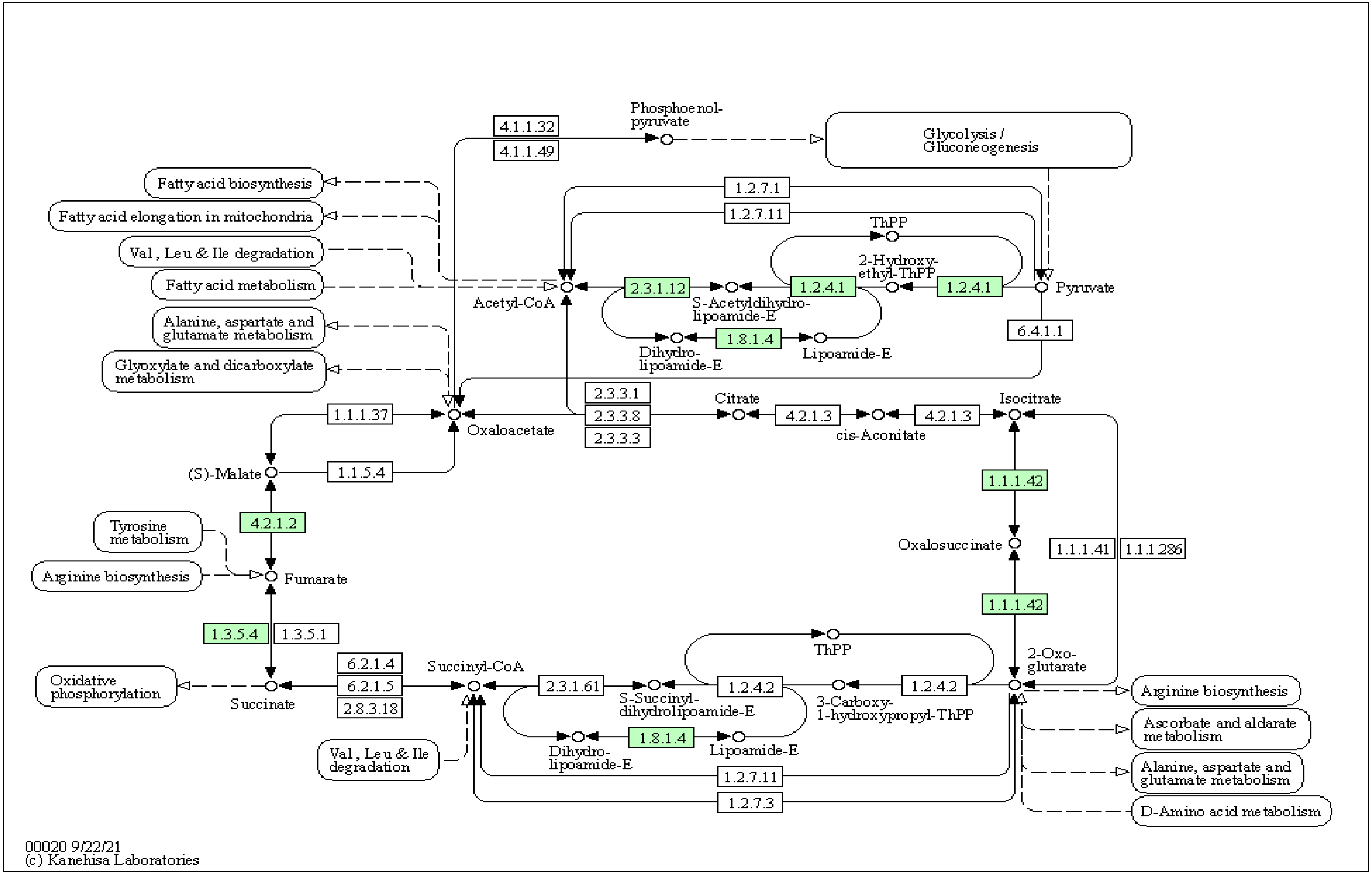


**Fig S3.** The graphical presentation of the **Tricarboxylic acid cycle (TCA)** of *Limosilactobacillus fermentum* AGA52 was obtained from KEGG Mapper (Green coloured EC number indicates the presence of the pathway enzymes).

**Table S10.** Comparison of carbohydrate fermentation patterns between *Limosilactobacillus fermentum* AGA52 and previously studied ATCC 14931.

| **Sugar** | **Strain** | |
|---|---|---|
| | **AGA 52** | **ATCC 14931*** |
| Control | - | - |
| Glycerol | - | - |
| Erythritol | - | - |
| D-Arabinose | - | - |
| L-Arabinose | - | - |
| D-Ribose | + | + |
| **D-Xylose** | + | - |
| L-Xylose | - | - |
| Adonitol | - | - |
| Methyl-βD-xylopyranoside | - | - |
| D-Galactose | + | + |
| D-Glucose | + | + |
| D-Fructose | + | + |
| D-Mannose | - | - |
| D-Sorbose | - | - |
| D-Rhamnose | - | - |
| Dulcitol | - | - |
| Inositol | - | - |
| D-Mannitol | - | - |
| D-Sorbitol | - | - |
| Methyl-αD-mannopyranoside | - | - |
| Methyl-αD-glucopyranoside | - | - |
| N-Acetylglucosamine | - | - |
| Amygdalin | - | - |
| Arbutin | - | - |
| **Esculin ferric citrate** | + | - |
| Salicin | - | - |
| **D-Cellobiose** | + | - |
| D-Maltose | + | + |
| D-Lactose | + | + |
| D-Melibiose | + | + |
| D-Sucrose | + | + |
| D-Trehalose | - | - |
| Inulin | - | - |
| D-Melezitose | - | - |
| D-Raffinose | + | + |
| Amidon (Starch) | - | - |
| Glycogen | - | - |
| Xylitol | - | - |
| Gentiobiose | - | - |
| D-Turanose | - | - |
| D-Lyxose | - | - |
| D-Tagatose | - | - |
| D-Fucose | - | - |
| L-Fucose | - | - |
| D-Arabitol | - | - |
| L-Arabitol | - | - |
| **Gluconate** | - | + |
| 2-Keto-gluconate | - | - |
| **5-Keto-gluconate** | + | - |

*ATCC 14931 studied by Buron Moles et al. 2019.

**Table S11.** KEGG (BlastKOALA) orthology search results for the enzymes responsible for carbohydrate metabolism.

| **Glycolysis / Gluconeogenesis** | | | | |
|---|---|---|---|---|
| **#** | **KEGG Entry** | **Symbol** | **Definition** | **Copy Number** |
| **1** | **K00016** | **LDH, ldh** | **L-lactate dehydrogenase [EC:1.1.1.27]** | **4** |
| 2 | K00134 | GAPDH, gapA | glyceraldehyde 3-phosphate dehydrogenase (phosphorylating) [EC:1.2.1.12] | 1 |
| 3 | K00161 | PDHA, pdhA | pyruvate dehydrogenase E1 component alpha subunit [EC:1.2.4.1] | 1 |
| 4 | K00162 | PDHB, pdhB | pyruvate dehydrogenase E1 component beta subunit [EC:1.2.4.1] | 1 |
| 5 | K00382 | DLD, lpd, pdhD | dihydrolipoamide dehydrogenase [EC:1.8.1.4] | 1 |
| 6 | K00627 | DLAT, aceF, pdhC | pyruvate dehydrogenase E2 component (dihydrolipoamide acetyltransferase) [EC:2.3.1.12] | 1 |
| 7 | K00873 | PK, pyk | pyruvate kinase [EC:2.7.1.40] | 1 |
| 8 | K00927 | PGK, pgk | phosphoglycerate kinase [EC:2.7.2.3] | 1 |
| 9 | K01223 | E3.2.1.86B, bglA | 6-phospho-beta-glucosidase [EC:3.2.1.86] | 1 |
| 10 | K01689 | ENO, eno | enolase [EC:4.2.1.11] | 1 |
| 11 | K01785 | galM, GALM | aldose 1-epimerase [EC:5.1.3.3] | 1 |
| 12 | K01803 | TPI, tpiA | triosephosphate isomerase (TIM) [EC:5.3.1.1] | 2 |
| **13** | **K01810** | **GPI, pgi** | **glucose-6-phosphate isomerase [EC:5.3.1.9]** | **3** |
| 14 | K01834 | PGAM, gpmA | 2,3-bisphosphoglycerate-dependent phosphoglycerate mutase [EC:5.4.2.11] | 2 |
| 15 | K01835 | pgm | phosphoglucomutase [EC:5.4.2.2] | 1 |
| 16 | K04072 | adhE | acetaldehyde dehydrogenase / alcohol dehydrogenase [EC:1.2.1.10 1.1.1.1] | 1 |
| 17 | K13953 | adhP | alcohol dehydrogenase, propanol-preferring [EC:1.1.1.1] | 2 |
| 18 | K25026 | glk | glucokinase [EC:2.7.1.2] | 1 |
| **Citrate cycle (TCA cycle)** | | | | |
| **#** | **KEGG Entry** | **Symbol** | **Definition** | **Copy Number** |
| 1 | K00031 | IDH1, IDH2, icd | isocitrate dehydrogenase [EC:1.1.1.42] | 1 |
| 2 | K00161 | PDHA, pdhA | pyruvate dehydrogenase E1 component alpha subunit [EC:1.2.4.1] | 1 |
| 3 | K00162 | PDHB, pdhB | pyruvate dehydrogenase E1 component beta subunit [EC:1.2.4.1] | 1 |
| 4 | K00244 | frdA | fumarate reductase flavoprotein subunit [EC:1.3.5.4] | 1 |
| 5 | K00382 | DLD, lpd, pdhD | dihydrolipoamide dehydrogenase [EC:1.8.1.4] | 1 |
| 6 | K00627 | DLAT, aceF, pdhC | pyruvate dehydrogenase E2 component (dihydrolipoamide acetyltransferase) [EC:2.3.1.12] | 1 |
| 7 | K01679 | E4.2.1.2B, fumC, FH | fumarate hydratase, class II [EC:4.2.1.2] | 1 |

**Table S11.** KEGG (BlastKOALA) orthology search results for the enzymes responsible for carbohydrate metabolism (continues).

**Pentose phosphate pathway**

| # | KEGG Entry | Symbol | Definition | Copy Number |
|---|---|---|---|---|
| 1 | K00033 | PGD, gnd, gntZ | 6-phosphogluconate dehydrogenase [EC:1.1.1.44 1.1.1.343] | 1 |
| 2 | K00036 | G6PD, zwf | glucose-6-phosphate 1-dehydrogenase [EC:1.1.1.49 1.1.1.363] | 1 |
| **3** | **K00615** | **E2.2.1.1, tktA, tktB** | **transketolase [EC:2.2.1.1]** | **2** |
| 4 | K00616 | E2.2.1.2, talA, talB | transaldolase [EC:2.2.1.2] | 1 |
| 5 | K00852 | rbsK, RBKS | ribokinase [EC:2.7.1.15] | 3 |
| 6 | K00948 | PRPS, prsA | ribose-phosphate pyrophosphokinase [EC:2.7.6.1] | 2 |
| 7 | K01619 | deoC, DERA | deoxyribose-phosphate aldolase [EC:4.1.2.4] | 1 |
| **8** | **K01621** | **xfp, xpk** | **xylulose-5-phosphate/fructose-6-phosphate phosphoketolase [EC:4.1.2.9/ 4.1.2.22]** | **1** |
| **9** | **K01783** | **rpe, RPE** | **ribulose-phosphate 3-epimerase [EC:5.1.3.1]** | **1** |
| 10 | K01807 | rpiA | ribose 5-phosphate isomerase A [EC:5.3.1.6] | 2 |
| **11** | **K01810** | **GPI, pgi** | **glucose-6-phosphate isomerase [EC:5.3.1.9]** | **3** |
| 12 | K01835 | pgm | phosphoglucomutase [EC:5.4.2.2] | 1 |
| 13 | K07404 | pgl | 6-phosphogluconolactonase [EC:3.1.1.31] | 2 |
| 14 | K25031 | gntK | gluconokinase [EC:2.7.1.12] | 1 |

**Pentose and glucuronate interconversions**

| # | KEGG Entry | Symbol | Definition | Copy Number |
|---|---|---|---|---|
| 1 | K00065 | kduD | 2-dehydro-3-deoxy-D-gluconate 5-dehydrogenase [EC:1.1.1.127] | 1 |
| 2 | K00854 | xylB, XYLB | xylulokinase [EC:2.7.1.17] | 1 |
| 3 | K00963 | UGP2, galU, galF | UTP--glucose-1-phosphate uridylyltransferase [EC:2.7.7.9] | 1 |
| 4 | K01783 | rpe, RPE | ribulose-phosphate 3-epimerase [EC:5.1.3.1] | 1 |
| **5** | **K01804** | **araA** | **L-arabinose isomerase [EC:5.3.1.4]** | **1** |
| 6 | K01805 | xylA | xylose isomerase [EC:5.3.1.5] | 1 |

**Table S11.** KEGG (BlastKOALA) orthology search results for the enzymes responsible for carbohydrate metabolism (continues).

| **Fructose and mannose metabolism** | | | | |
|---|---|---|---|---|
| **#** | **KEGG Entry** | **Symbol** | **Definition** | **Copy Number** |
| 1 | K00847 | E2.7.1.4, scrK | fructokinase [EC:2.7.1.4] | 1 |
| 2 | K01803 | TPI, tpiA | triosephosphate isomerase (TIM) [EC:5.3.1.1] | 2 |
| 3 | K01805 | xylA | xylose isomerase [EC:5.3.1.5] | 1 |
| 4 | K01809 | manA, MPI | mannose-6-phosphate isomerase [EC:5.3.1.8] | 1 |
| 5 | K02768 | fruB | fructose PTS system EIIA component [EC:2.7.1.202] | 1 |
| 6 | K02769 | fruAb | fructose PTS system EIIB component [EC:2.7.1.202] | 1 |
| 7 | K02770 | fruA | fructose PTS system EIIBC or EIIC component [EC:2.7.1.202] | 1 |
| 8 | K02794 | manX | mannose PTS system EIIAB component [EC:2.7.1.191] | 1 |
| 9 | K02795 | manY | mannose PTS system EIIC component | 1 |
| 10 | K02796 | manZ | mannose PTS system EIID component | 1 |
| 11 | K17195 | alsE | D-allulose-6-phosphate 3-epimerase [EC:5.1.3.-] | 1 |

| **Galactose metabolism** | | | | |
|---|---|---|---|---|
| **#** | **KEGG Entry** | **Symbol** | **Definition** | **Copy Number** |
| **1** | **K00849** | **galK** | **galactokinase [EC:2.7.1.6]** | **1** |
| **2** | **K00963** | **UGP2, galU, galF** | **UTP--glucose-1-phosphate uridylyltransferase [EC:2.7.7.9]** | **1** |
| 3 | K00965 | galT, GALT | UDPglucose--hexose-1-phosphate uridylyltransferase [EC:2.7.7.12] | 1 |
| 4 | K01182 | IMA, malL | oligo-1,6-glucosidase [EC:3.2.1.10] | 1 |
| **5** | **K01190** | **lacZ** | **beta-galactosidase [EC:3.2.1.23]** | **2** |
| 6 | K01193 | INV, sacA | beta-fructofuranosidase [EC:3.2.1.26] | 1 |
| **7** | **K01784** | **galE, GALE** | **UDP-glucose 4-epimerase [EC:5.1.3.2]** | **2** |
| **8** | **K01785** | **galM, GALM** | **aldose 1-epimerase [EC:5.1.3.3]** | **1** |
| 9 | K01835 | pgm | phosphoglucomutase [EC:5.4.2.2] | 1 |
| 10 | K01854 | glf | UDP-galactopyranose mutase [EC:5.4.99.9] | 1 |
| 11 | K02775 | gatC, sgcC | galactitol PTS system EIIC component | 1 |
| 12 | K07407 | E3.2.1.22B, galA, rafA | alpha-galactosidase [EC:3.2.1.22] | 1 |
| 13 | K25026 | glk | glucokinase [EC:2.7.1.2] | 1 |

**Table S11.** KEGG (BlastKOALA) orthology search results for the enzymes responsible for carbohydrate metabolism (continues).

**Ascorbate and aldarate metabolism**

| # | KEGG Entry | Symbol | Definition | Copy Number |
|---|---|---|---|---|
| 1 | K02821 | ulaC, sgaA | ascorbate PTS system EIIA or EIIAB component [EC:2.7.1.194] | 1 |
| 2 | K02822 | ulaB, sgaB | ascorbate PTS system EIIB component [EC:2.7.1.194] | 1 |
| 3 | K03475 | ulaA, sgaT | ascorbate PTS system EIIC component | 1 |

**Starch and sucrose metabolism**

| # | KEGG Entry | Symbol | Definition | Copy Number |
|---|---|---|---|---|
| 1 | K00691 | mapA | maltose phosphorylase [EC:2.4.1.8] | 1 |
| 2 | K00847 | E2.7.1.4, scrK | fructokinase [EC:2.7.1.4] | 1 |
| 3 | K00963 | UGP2, galU, galF | UTP--glucose-1-phosphate uridylyltransferase [EC:2.7.7.9] | 1 |
| 4 | K01182 | IMA, malL | oligo-1,6-glucosidase [EC:3.2.1.10] | 1 |
| 5 | K01193 | INV, sacA | beta-fructofuranosidase [EC:3.2.1.26] | 1 |
| 6 | K01223 | E3.2.1.86B, bglA | 6-phospho-beta-glucosidase [EC:3.2.1.86] | 1 |
| **7** | **K01810** | **GPI, pgi** | **glucose-6-phosphate isomerase [EC:5.3.1.9]** | **3** |
| 8 | K01835 | pgm | phosphoglucomutase [EC:5.4.2.2] | 1 |
| 9 | K01838 | pgmB | beta-phosphoglucomutase [EC:5.4.2.6] | 1 |
| 10 | K02761 | celB, chbC | cellobiose PTS system EIIC component | 1 |
| 11 | K02810 | scrA, sacP, sacX, ptsS | sucrose PTS system EIIBCA or EIIBC component [EC:2.7.1.211] | 1 |
| 12 | K20811 | inuJ | inulosucrase [EC:2.4.1.9] | 1 |
| 13 | K25026 | glk | glucokinase [EC:2.7.1.2] | 1 |

**Table S11.** KEGG (BlastKOALA) orthology search results for the enzymes responsible for carbohydrate metabolism (continues).

| **Amino sugar and nucleotide sugar metabolism** | | | | |
|---|---|---|---|---|
| **#** | **KEGG Entry** | **Symbol** | **Definition** | **Copy Number** |
| 1 | K00075 | murB | UDP-N-acetylmuramate dehydrogenase [EC:1.3.1.98] | 1 |
| 2 | K00790 | murA | UDP-N-acetylglucosamine 1-carboxy vinyl transferase [EC:2.5.1.7] | 2 |
| 3 | K00820 | glmS, GFPT | glutamine-fructose-6-phosphate transaminase (isomerizing) [EC:2.6.1.16] | 1 |
| 4 | K00847 | E2.7.1.4, scrK | fructokinase [EC:2.7.1.4] | 1 |
| 5 | K00849 | galK | galactokinase [EC:2.7.1.6] | 1 |
| 6 | K00963 | UGP2, galU, galF | UTP--glucose-1-phosphate uridylyltransferase [EC:2.7.7.9] | 1 |
| 7 | K00965 | galT, GALT | UDP-glucose-hexose-1-phosphate uridylyltransferase [EC:2.7.7.12] | 1 |
| 8 | K01209 | abfA | alpha-L-arabinofuranosidase [EC:3.2.1.55] | 1 |
| 9 | K01784 | galE, GALE | UDP-glucose 4-epimerase [EC:5.1.3.2] | 2 |
| 10 | K01809 | manA, MPI | mannose-6-phosphate isomerase [EC:5.3.1.8] | 1 |
| **11** | **K01810** | **GPI, pgi** | **glucose-6-phosphate isomerase [EC:5.3.1.9]** | **3** |
| 12 | K01835 | pgm | phosphoglucomutase [EC:5.4.2.2] | 1 |
| 13 | K01854 | glf | UDP-galactopyranose mutase [EC:5.4.99.9] | 1 |
| 14 | K02794 | manX | mannose PTS system EIIAB component [EC:2.7.1.191] | 1 |
| 15 | K02795 | manY | mannose PTS system EIIC component | 1 |
| 16 | K02796 | manZ | mannose PTS system EIID component | 1 |
| 17 | K03431 | glmM | phosphoglucosamine mutase [EC:5.4.2.10] | 1 |
| 18 | K04042 | glmU | bifunctional UDP-N-acetylglucosamine pyrophosphorylase / glucosamine-1-phosphate N-acetyltransferase [EC:2.7.7.23 2.3.1.157] | 1 |
| 19 | K25026 | glk | glucokinase [EC:2.7.1.2] | 1 |

**Table S11.** KEGG (BlastKOALA) orthology search results for the enzymes responsible for carbohydrate metabolism (continues).

| **Pyruvate metabolism** | | | | |
|---|---|---|---|---|
| **#** | **KEGG Entry** | **Symbol** | **Definition** | **Copy Number** |
| **1** | **K00016** | **LDH, ldh** | **L-lactate dehydrogenase [EC:1.1.1.27]** | **4** |
| 2 | K00161 | PDHA, pdhA | pyruvate dehydrogenase E1 component alpha subunit [EC:1.2.4.1] | 1 |
| 3 | K00162 | PDHB, pdhB | pyruvate dehydrogenase E1 component beta subunit [EC:1.2.4.1] | 1 |
| 4 | K00244 | frdA | fumarate reductase flavoprotein subunit [EC:1.3.5.4] | 1 |
| 5 | K00382 | DLD, lpd, pdhD | dihydrolipoamide dehydrogenase [EC:1.8.1.4] | 1 |
| 6 | K00625 | E2.3.1.8, pta | phosphate acetyltransferase [EC:2.3.1.8] | 1 |
| 7 | K00626 | ACAT, atoB | acetyl-CoA C-acetyltransferase [EC:2.3.1.9] | 1 |
| 8 | K00627 | DLAT, aceF, pdhC | pyruvate dehydrogenase E2 component (dihydrolipoamide acetyltransferase) [EC:2.3.1.12] | 1 |
| 9 | K00873 | PK, pyk | pyruvate kinase [EC:2.7.1.40] | 1 |
| 10 | K00925 | ackA | acetate kinase [EC:2.7.2.1] | 1 |
| 11 | K01512 | acyP | acylphosphatase [EC:3.6.1.7] | 1 |
| 12 | K01679 | E4.2.1.2B, fumC, FH | fumarate hydratase, class II [EC:4.2.1.2] | 1 |
| 13 | K01759 | GLO1, gloA | lactoylglutathione lyase [EC:4.4.1.5] | 1 |
| 14 | K01961 | accC | acetyl-CoA carboxylase, biotin carboxylase subunit [EC:6.4.1.2 6.3.4.14] | 2 |
| 15 | K01962 | accA | acetyl-CoA carboxylase carboxyl transferase subunit alpha [EC:6.4.1.2 2.1.3.15] | 1 |
| 16 | K01963 | accD | acetyl-CoA carboxylase carboxyl transferase subunit beta [EC:6.4.1.2 2.1.3.15] | 1 |
| 17 | K02160 | accB, bccP | acetyl-CoA carboxylase biotin carboxyl carrier protein | 2 |
| **18** | **K03777** | **dld** | **D-lactate dehydrogenase (quinone) [EC:1.1.5.12]** | **1** |
| **19** | **K03778** | **ldhA** | **D-lactate dehydrogenase [EC:1.1.1.28]** | **3** |
| **20** | **K04072** | **adhE** | **acetaldehyde dehydrogenase / alcohol dehydrogenase [EC:1.2.1.10 1.1.1.1]** | **1** |
| **21** | **K13953** | **adhP** | **alcohol dehydrogenase, propanol-preferring [EC:1.1.1.1]** | **2** |
| 22 | K22212 | mleA, mleS | malolactic enzyme [EC:4.1.1.101] | 1 |

**Table S11.** KEGG (BlastKOALA) orthology search results for the enzymes responsible for carbohydrate metabolism (continues).

**Glyoxylate and dicarboxylate metabolism**

| # | KEGG Entry | Symbol | Definition | Copy Number |
|---|---|---|---|---|
| 1 | K00382 | DLD, lpd, pdhD | dihydrolipoamide dehydrogenase [EC:1.8.1.4] | 1 |
| 2 | K00600 | glyA, SHMT | glycine hydroxymethyltransferase [EC:2.1.2.1] | 1 |
| 3 | K00626 | ACAT, atoB | acetyl-CoA C-acetyltransferase [EC:2.3.1.9] | 1 |
| 4 | K00865 | glxK, garK | glycerate 2-kinase [EC:2.7.1.165] | 1 |
| 5 | K01915 | glnA, GLUL | glutamine synthetase [EC:6.3.1.2] | 1 |
| 6 | K02437 | gcvH, GCSH | glycine cleavage system H protein | 1 |

**Propanoate metabolism**

| # | KEGG Entry | Symbol | Definition | Copy Number |
|---|---|---|---|---|
| 1 | K00005 | gldA | glycerol dehydrogenase [EC:1.1.1.6] | 1 |
| **2** | **K00016** | **LDH, ldh** | **L-lactate dehydrogenase [EC:1.1.1.27]** | **4** |
| 3 | K00086 | dhaT | 1,3-propanediol dehydrogenase [EC:1.1.1.202] | 1 |
| 4 | K00382 | DLD, lpd, pdhD | dihydrolipoamide dehydrogenase [EC:1.8.1.4] | 1 |
| 5 | K00625 | E2.3.1.8, pta | phosphate acetyltransferase [EC:2.3.1.8] | 1 |
| 6 | K00925 | ackA | acetate kinase [EC:2.7.2.1] | 1 |
| 7 | K01961 | accC | acetyl-CoA carboxylase, biotin carboxylase subunit [EC:6.4.1.2 6.3.4.14] | 2 |
| 8 | K01962 | accA | acetyl-CoA carboxylase carboxyl transferase subunit alpha [EC:6.4.1.2 2.1.3.15] | 1 |
| 9 | K01963 | accD | acetyl-CoA carboxylase carboxyl transferase subunit beta [EC:6.4.1.2 2.1.3.15] | 1 |
| 10 | K02160 | accB, bccP | acetyl-CoA carboxylase biotin carboxyl carrier protein | 2 |

**C5-Branched dibasic acid metabolism**

| # | KEGG Entry | Symbol | Definition | Copy Number |
|---|---|---|---|---|
| 1 | K00052 | leuB, IMDH | 3-isopropylmalate dehydrogenase [EC:1.1.1.85] | 1 |
| 2 | K01575 | alsD, budA, aldC | acetolactate decarboxylase [EC:4.1.1.5] | 1 |
| 3 | K01652 | E2.2.1.6L, ilvB, ilvG, ilvI | acetolactate synthase I/II/III large subunit [EC:2.2.1.6] | 1 |
| 4 | K01703 | leuC, IPMI-L | 3-isopropylmalate/(R)-2-methylmalate dehydratase large subunit [EC:4.2.1.33 4.2.1.35] | 1 |
| 5 | K01704 | leuD, IPMI-S | 3-isopropylmalate/(R)-2-methylmalate dehydratase small subunit [EC:4.2.1.33 4.2.1.35] | 1 |

**Table S11.** KEGG (BlastKOALA) orthology search results for the enzymes responsible for carbohydrate metabolism (continues).

**Inositol phosphate metabolism**

| # | KEGG Entry | Symbol | Definition | Copy Number |
|---|---|---|---|---|
| 1 | K01092 | E3.1.3.25, IMPA, suhB | myo-inositol-1(or 4)-monophosphatase [EC:3.1.3.25] | 1 |
| 2 | K01803 | TPI, tpiA | triosephosphate isomerase (TIM) [EC:5.3.1.1] | 2 |
| 3 | K22230 | iolU | scyllo-inositol 2-dehydrogenase (NADP+) [EC:1.1.1.-] | 1 |

**Butanoate metabolism**

| # | KEGG Entry | Symbol | Definition | Copy Number |
|---|---|---|---|---|
| 1 | K00004 | BDH, butB | (R,R)-butanediol dehydrogenase / meso-butanediol dehydrogenase / diacetyl reductase [EC:1.1.1.4 1.1.1.- 1.1.1.303] | 1 |
| 2 | K00022 | HADH | 3-hydroxyacyl-CoA dehydrogenase [EC:1.1.1.35] | 1 |
| 3 | K00135 | gabD | succinate-semialdehyde dehydrogenase / glutarate-semialdehyde dehydrogenase [EC:1.2.1.16 1.2.1.79 1.2.1.20] | 1 |
| 4 | K00244 | frdA | fumarate reductase flavoprotein subunit [EC:1.3.5.4] | 1 |
| 5 | K00626 | ACAT, atoB | acetyl-CoA C-acetyltransferase [EC:2.3.1.9] | 1 |
| 6 | K01575 | alsD, budA, aldC | acetolactate decarboxylase [EC:4.1.1.5] | 1 |
| 7 | K01580 | E4.1.1.15, gadB, gadA, GAD | glutamate decarboxylase [EC:4.1.1.15] | 1 |
| 8 | K01641 | HMGCS | hydroxymethylglutaryl-CoA synthase [EC:2.3.3.10] | 1 |
| 9 | K01652 | E2.2.1.6L, ilvB, ilvG, ilvI | acetolactate synthase I/II/III large subunit [EC:2.2.1.6] | 1 |
| 10 | K03366 | butA, budC | meso-butanediol dehydrogenase / (S,S)-butanediol dehydrogenase / diacetyl reductase [EC:1.1.1.- 1.1.1.76 1.1.1.304] | 1 |
| 11 | K04072 | adhE | acetaldehyde dehydrogenase / alcohol dehydrogenase [EC:1.2.1.10 1.1.1.1] | 1 |
| 12 | K18009 | budC | meso-butanediol dehydrogenase / (S,S)-butanediol dehydrogenase / diacetyl reductase [EC:1.1.1.- 1.1.1.76 1.1.1.304] | 1 |
| 13 | K23259 | adh | isopropanol dehydrogenase (NADP+) [EC:1.1.1.80] | 1 |

**Table S12.** KEGG (BlastKOALA) orthology search results for ABC transporters.

**ABC Transporters**

| # | KEGG Entry | Symbol | Definition | Copy Number |
|---|---|---|---|---|
| 1 | K02036 | pstB | phosphate transport system ATP-binding protein [EC:7.3.2.1] | 2 |
| 2 | K02037 | pstC | phosphate transport system permease protein | 2 |
| 3 | K02038 | pstA | phosphate transport system permease protein | 2 |
| 4 | K02040 | pstS | phosphate transport system substrate-binding protein | 2 |
| 5 | K02071 | metN | D-methionine transport system ATP-binding protein | 1 |
| 6 | K02072 | metI | D-methionine transport system permease protein | 1 |
| 7 | K02073 | metQ | D-methionine transport system substrate-binding protein | 3 |
| 8 | K02424 | fliY, tcyA | L-cystine transport system substrate-binding protein | 2 |
| 9 | K03523 | bioY | biotin transport system substrate-specific component | 1 |
| 10 | K05845 | opuC | osmoprotectant transport system substrate-binding protein | 1 |
| 11 | K05846 | opuBD | osmoprotectant transport system permease protein | 2 |
| 12 | K05847 | opuA | osmoprotectant transport system ATP-binding protein [EC:7.6.2.9] | 1 |
| 13 | K06726 | rbsD | D-ribose pyranase [EC:5.4.99.62] | 1 |
| 14 | K10009 | tcyB, yecS | L-cystine transport system permease protein | 2 |
| 15 | K10010 | tcyC, yecC | L-cystine transport system ATP-binding protein [EC:7.4.2.1] | 1 |
| 16 | K10036 | glnH | glutamine transport system substrate-binding protein | 1 |
| 17 | K10037 | glnP | glutamine transport system permease protein | 1 |
| 18 | K10038 | glnQ | glutamine transport system ATP-binding protein [EC:7.4.2.1] | 1 |
| 19 | K16012 | cydC | ATP-binding cassette, subfamily C, bacterial CydC | 1 |
| 20 | K16013 | cydD | ATP-binding cassette, subfamily C, bacterial CydD | 1 |

**Table S12.** KEGG (BlastKOALA) orthology search results for ABC transporters.

| ABC Transporters | | | | |
|---|---|---|---|---|
| **#** | **KEGG Entry** | **Symbol** | **Definition** | **Copy Number** |
| 21 | K16785 | ecfT | energy-coupling factor transport system permease protein | 2 |
| 22 | K16786 | ecfA1 | energy-coupling factor transport system ATP-binding protein [EC:7.-.-.-] | 1 |
| 23 | K16787 | ecfA2 | energy-coupling factor transport system ATP-binding protein [EC:7.-.-.-] | 1 |
| 24 | K16961 | yxeM | putative S-methyl cysteine transport system substrate-binding protein | 1 |
| 25 | K16962 | yxeN | putative S-methyl cysteine transport system permease protein | 1 |
| 26 | K16963 | yxeO | putative S-methyl cysteine transport system ATP-binding protein | 1 |
| 27 | K17077 | artQ | arginine/lysine/histidine transport system permease protein | 1 |
| **28** | **K18887** | **efrA, efrE** | **ATP-binding cassette, subfamily B, multidrug efflux pump** | **1** |
| **29** | **K18888** | **efrB, efrF** | **ATP-binding cassette, subfamily B, multidrug efflux pump** | **1** |
| 30 | K23059 | artP, artI | arginine/lysine/histidine transporter system substrate-binding protein | 2 |
| 31 | K23060 | artR, artM | arginine/lysine/histidine transport system ATP-binding protein [EC:7.4.2.1] | 1 |

**Table S13.** KEGG (BlastKOALA) orthology search results for phosphotransferase system (PTS).

| **Phosphotransferase system (PTS)** | | | | |
|---|---|---|---|---|
| **#** | **KEGG Entry** | **Symbol** | **Definition** | **Copy Number** |
| 1 | K02757 | bglF, bglP | beta-glucoside PTS system EIICBA component [EC:2.7.1.-] | 1 |
| 2 | K02761 | celB, chbC | cellobiose PTS system EIIC component | 1 |
| 3 | K02768 | fruB | fructose PTS system EIIA component [EC:2.7.1.202] | 1 |
| 4 | K02769 | fruAb | fructose PTS system EIIB component [EC:2.7.1.202] | 1 |
| 5 | K02770 | fruA | fructose PTS system EIIBC or EIIC component [EC:2.7.1.202] | 1 |
| 6 | K02775 | gatC, sgcC | galactitol PTS system EIIC component | 1 |
| **7** | **K02784** | **ptsH** | **phosphocarrier protein HPr** | **1** |
| 8 | K02794 | manX | mannose PTS system EIIAB component [EC:2.7.1.191] | 1 |
| 9 | K02795 | manY | mannose PTS system EIIC component | 1 |
| 10 | K02796 | manZ | mannose PTS system EIID component | 1 |
| 11 | K02810 | scrA, sacP, sacX, ptsS | sucrose PTS system EIIBCA or EIIBC component [EC:2.7.1.211] | 1 |
| 12 | K02821 | ulaC, sgaA | ascorbate PTS system EIIA or EIIAB component [EC:2.7.1.194] | 1 |
| 13 | K02822 | ulaB, sgaB | ascorbate PTS system EIIB component [EC:2.7.1.194] | 1 |
| 14 | K03475 | ulaA, sgaT | ascorbate PTS system EIIC component | 1 |
| **15** | **K08483** | **ptsI** | **phosphoenolpyruvate-protein phosphotransferase (PTS system enzyme I) [EC:2.7.3.9]** | **2** |

**Table S14.** Comparison of presence or absence of the key enzymes responsible for EMP (Embden–Meyerhof–Parnas), PK (phosphoketolase) and Leloir pathways via RASTtk by descending order.

| Enzyme | E.C. number | Gene copy number of the strains | | | | | | | |
|---|---|---|---|---|---|---|---|---|---|
| | | AGA52 | DSM 20052 | ATCC 14931 | SK152 | YLF016 | B44 | IFO3956 | CECT5716 |
| **6-phosphofructokinase 1** | 2.7.1.11 | 0 | 0 | 0 | 0 | 0 | 0 | 0 | 0 |
| **Fructose-bisphosphate aldolase** | 4.1.2.13 | 0 | 0 | 0 | 0 | 0 | 0 | 0 | 0 |
| Glucose-6-phosphate isomerase | 5.3.1.9 | 6 | 2 | 1 | 2 | 2 | 2 | 2 | 5 |
| Transketolase | 2.2.1.1 | 2 | 2 | 2 | 2 | 2 | 2 | 2 | 2 |
| Phosphoketolase | 4.1.2.9/4.1.2.22 | 1 | 1 | 1 | 1 | 1 | 1 | 1 | 1 |
| L-arabinose isomerase | 5.3.1.4 | 1 | 1 | 1 | 1 | 0 | 1 | 1 | 1 |
| **L-ribulose kinase** | 2.7.1.19 | 0 | 0 | 0 | 0 | 0 | 0 | 0 | 0 |
| Ribulose-phosphate 3-epimerase | 5.1.3.1 | 2 | 1 | 1 | 2 | 1 | 1 | 1 | 1 |
| galactokinase | 2.7.1.6 | 1 | 1 | 1 | 1 | 1 | 1 | 1 | 1 |
| UTP--glucose-1-phosphate uridylyltransferase | 2.7.7.9 | 1 | 1 | 1 | 1 | 1 | 1 | 1 | 1 |
| UDP-glucose 4-epimerase | 5.1.3.2 | 2 | 2 | 1 | 2 | 1 | 1 | 1 | 1 |
| Aldose 1-epimerase | 5.1.3.3 | 1 | 1 | 1 | 1 | 1 | 1 | 1 | 1 |
| Beta-galactosidase | 3.2.1.23 | 2 | 2 | 2 | 2 | 2 | 2 | 3 | 1 |

**Table S15.** Putative probiotic and psychobiotic function-related genes are found in the genome of *Limosilactobacillus fermentum* AGA52 (CP091132.1).

| Gene | Putative Function | Response | Origin | Location | Strand |
|---|---|---|---|---|---|
| **Stress resistance genes** | | | | | |
| ***dltD*** (D-alanyl-lipoteichoic acid biosynthesis protein) | d- anylation of LTA | Acid and defensin Resistance | *L. fermentum* | 1643923-1645215 | + |
| ***dltA*** (D-alanine--poly(phosphoribitol) ligase subunit) | d- anylation of LTA | Acid and defensin Resistance | *L. fermentum* | 1640888-1642417 | + |
| ***dltB,*** D-alanyl-lipoteichoic acid biosynthesis protein | d- anylation of LTA | Acid and defensin Resistance | *L. fermentum* | 1642417-1643643 | + |
| ***xylA*** **(**Xylose isomerase**)** | Interconversion of D-xylose and D-xylulose | Gut persistence | *L. fermentum* | 723929-725278 | - |
| ***NapA***, **L1970_03860**, Na+/H+ antiporter | Cation:proton antiporter | Acid resistance | *L. fermentum* | 791603-792766 | + |
| ***NhaC***, Na+/H+ antiporter | Cation:proton antiporter | Acid resistance | *L. fermentum* | 997161-998558 | + |
| **L1970_07245,** ***NhaP-type*** Na+/H+ and K+/H+ antiporter | Cation:proton antiporter | Acid resistance | *L. fermentum* | 1467327-1469288 | - |
| ***YbiT***, **L1970_04860**, Bis-ABC ATPase, | ATP-binding cassette domain-containing protein | Acid resistance | *L. fermentum* | 983519-985144 | - |
| ***Uup***, Bis-ABC ATPase **L1970_00870**, | ABC-F family ATP-binding cassette domain-containing protein | Acid resistance | *L. fermentum* | 170046-171932 | + |
| ***YheS***, Bis-ABC ATPase, | ABC-F type ribosomal protection protein | Acid resistance | *L. fermentum* | 268068-270011 | + |
| **L1970_06450** | cation-transport ATPase, E1-E2 family | Bile salt resistance | *L. fermentum* | 1305342-1305797 | - |
| **L1970_07400**, Bis-ABC ATPase **SPy1206** | ATP-binding cassette domain-containing protein | Acid resistance | *L. fermentum* | 1499925-1501469 | - |
| **L1970_07400**, ATP synthase epsilon chain (EC 3.6.3.14) | F0F1-type ATP synthase | Acid resistance | *L. fermentum* | 1499925-1501469 | - |
| ***AtpD***, ATP synthase beta chain (EC 3.6.3.14) | F0F1-type ATP synthase | Acid resistance | *L. fermentum* | 1326339-1327760 | - |
| **L1970_06575,** ATP synthase gamma chain (EC 3.6.3.14) | F0F1-type ATP synthase | Acid resistance | *L. fermentum* | 1327783-1328718 | - |
| ***AtpA***, ATP synthase alpha chain (EC 3.6.3.14) | F0F1-type ATP synthase | Acid resistance | *L. fermentum* | 1328750-1330288 | - |
| **L1970_06585**, ATP synthase delta chain (EC 3.6.3.14) | F0F1-type ATP synthase | Acid resistance | *L. fermentum* | 1330316-1330861 | - |
| ***AtpF***, ATP synthase F0 sector subunit b (EC 3.6.3.14) | F0F1-type ATP synthase | Acid resistance | *L. fermentum* | 1330854-1331360 | - |
| ***AtpE***, ATP synthase F0 sector subunit c (EC 3.6.3.14) | F0F1-type ATP synthase | Acid resistance | *L. fermentum* | 1331406-1331618 | - |
| ***AtpB***, ATP synthase F0 sector subunit a (EC 3.6.3.14) | F0F1-type ATP synthase | Acid resistance | *L. fermentum* | 1331647-1332357 | - |
| **Immunomodulation** | | | | | |
| ***DltB*** (D-alanyl transfer protein) | d- anylation of LTA | Anti-inflammatory potential in vitro in PBMCs and in vivo in a murine model of colitis | *L. fermentum* | 1642417-1643643 | + |
| **Anti-pathogenic effect** | | | | | |
| ***LuxS*** (S-ribosylhomocysteine lyase (EC 4.4.1.21), Autoinducer-2 production protein) | Autoinducer-2 production | Autoinduction ability | *L. fermentum* | 1607587-1608063 | - |

**Table S15.** Putative probiotic and psychobiotic function-related genes are found in the genome of *Limosilactobacillus fermentum* AGA52 (CP091132.1).

| Gene | Putative Function | Response | Origin | Location | Strand |
|---|---|---|---|---|---|
| **Exopolysaccharide biosynthesis responsible genes** | | | | | |
| **L1970_04765** Undecaprenyl-phosphate galactosephosphotransferase (EC 2.7.8.6) | Sugar transferase | Adhesion ability | *L. fermentum* | 996419-967072 | - |
| ***EpsC* L1970_08410** Tyrosine-protein kinase transmembrane modulator | Wzz/FepE/Etk N-terminal domain-containing protein | Adhesion ability | *L. fermentum* | 1713425-1714195 | + |
| ***EpsD*, L1970_08415** Tyrosine-protein kinase (EC 2.7.10.2) | CpsD/CapB family tyrosine-protein kinase | Adhesion ability | *L. fermentum* | 1714212-1714955 | + |
| **L1970_08420**, Manganese-dependent protein-tyrosine phosphatase (EC 3.1.3.48) | Exopolysaccharide biosynthesis protein | Adhesion ability | *L. fermentum* | 1714976-1715746 | + |
| **Lipoteichoic acid (LTA) synthesis responsible genes** | | | | | |
| ***MurJ*, Flippase, L1970_04740** | Type IV ATPase | Adhesion ability | *L. fermentum* | 961163-962581 | - |
| ***LtaS* Type IIb, L1970_07305**, Lipoteichoic acid synthase | LTA synthase family protein | Adhesion ability | *L. fermentum* | 1479646-1481775 | - |
| ***LafC*, L1970_08010**, Integral membrane protein | Accessory function in glycolipid and LTA synthesis | Adhesion ability | *L. fermentum* | 1623734-1624753 | - |
| ***LafB*, L1970_08015** Glycosyltransferase | Formation of Gal-Glc-DAG | Adhesion ability | *L. fermentum* | 1624755-1625780 | - |
| ***LafA*, L1970_08020** Glycosyltransferase | Formation of Glc-DAG | Adhesion ability | *L. fermentum* | 1625773-1626990 | - |
| ***dltD*** (D-alanyl-lipoteichoic acid biosynthesis protein) | d- anylation of LTA | Adhesion ability | *L. fermentum* | 1643923-1645215 | + |
| ***dltA*** (D-alanine--poly(phosphoribitol) ligase subunit) | d- anylation of LTA | Adhesion ability | *L. fermentum* | 1640888-1642417 | + |
| ***dltB*,** D-alanyl-lipoteichoic acid biosynthesis protein | d- anylation of LTA | Adhesion ability | *L. fermentum* | 1642417-1643643 | + |
| **Putative antioxidant activity responsible genes** | | | | | |
| ***TrxA*, L1970_00170** thioredoxin | Catalyzing electron flux from nicotinamide adenine dinucleotide phosphate | Antioxidant activity | *L. fermentum* | 31766-32083 | + |
| ***TrxA*,** *thioredoxin* | As same as mentioned above | Antioxidant activity | *L. fermentum* | 383142-383456 | - |
| ***Tpx***, Thiol peroxidase, Tpx-type (EC 1.11.1.15) | Alcohol and water formation from organic hydroperoxides | Antioxidant activity | *L. fermentum* | 421433-421927 | - |
| ***DsbA*,** bacterial thiol disulfide oxidoreductase | Catalyzes intrachain disulfide bond formation | Antioxidant activity | *L. fermentum* | 441969-442595 | - |
| ***TrxB***, Thioredoxin reductase (EC 1.8.1.9) | Pyrimidine conversions, NAD(P)/FAD-dependent oxidoreductase | Antioxidant activity | *L. fermentum* | 1474169-1475161 | + |
| ***TrxA*,** *thioredoxin* | Catalyzing electron flux from nicotinamide adenine dinucleotide phosphate | Antioxidant activity | *L. fermentum* | 1488228-1488548 | + |
| ***TrxB***, Thioredoxin reductase (EC 1.8.1.9) | Pyrimidine conversions, NAD(P)/FAD-dependent oxidoreductase | Antioxidant activity | *L. fermentum* | 1530130-1531071 | + |

**Table S15.** Putative probiotic and psychobiotic function-related genes are found in the genome of *Limosilactobacillus fermentum* AGA52 (CP091132.1).

| Gene | Putative Function | Response | Origin | Location | Strand |
|---|---|---|---|---|---|
| **Putative antioxidant activity responsible genes** | | | | | |
| ***ArsC***, Arsenate reductase (EC 1.20.4.4) thioredoxin-coupled, LMWP family | Prodution of non-toxic organoarsenical compounds | Antioxidant activity | *L. fermentum* | 1561836-1562252 | - |
| ***TrxA*,** thioredoxin | Catalyzing electron flux from nicotinamide adenine dinucleotide phosphate | Antioxidant activity | *L. fermentum* | 1946030-1946344 | + |
| ***Gor*, L1970_09685**, Putative glutathione reductase | Cellular control of reactive oxygen species | Antioxidant activity | *L. fermentum* | 1946498-1947829 | + |
| ***nrdH*,** putative glutaredoxin | Reduction of ribonucleotide reductase class Ib | Antioxidant activity | *L. fermentum* | 307953-308174 | + |
| **L1970_02430**, Arsenate reductase related protein, glutaredoxin family | Spx/MgsR family RNA polymerase-binding regulatory protein | Antioxidant activity | *L. fermentum* | 479278-479706 | - |
| **L1970_06355**, NADH dehydrogenase (EC 1.6.99.3) | NAD(P)/FAD-dependent oxidoreductase | Antioxidant activity | *L. fermentum* | 1286111-1287334 | + |
| **DNA and protein protection and repair** | | | | | |
| ***MsrB*** (peptide-methionine (R)-S-oxide reductase) | Methionine sulfoxide reductase | Persistence capacity *in vivo* | *L. fermentum* | 193260-193685 | + |
| ***ClpX*** (ATP-dependent Clp protease ATP-binding subunit) | Clp ATPase (chaperone) | Acid and bile tolerance | *L. fermentum* | 502507-503757 | + |
| ***ClpP*** (ATP-dependent Clp protease proteolytic subunit) | Clp ATPase (chaperone) | Persistence capacity *in vivo* | *L. fermentum* | 773713-774303 | + |
| ***ClpC*** (ATP-dependent Clp protease, ATP-binding subunit) | Clp ATPase (chaperone) | Acid and bile tolerance | *L. fermentum* | 1120468-1122972 | - |
| ***ClpE*** (ATP-dependent Clp protease, ATP-binding subunit) | Clp ATPase (chaperone) | Acid and bile tolerance | *L. fermentum* | 1629750-1631996 | + |
| **phage*ClpP*** (ATP-dependent Clp protease proteolytic subunit, L1970_04120) | Prophage Clp protease-like protein | Persistence capacity *in vivo* | *L. reuterii* | 849351-850022 | - |
| **Active removal of stressors** | | | | | |
| ***gadC*** (glutamate: GABA symporter) | GABA transporter | Acid tolerance | *L. fermentum* | 1080245-1081537 | - |
| ***cbh*** (Choloylglycine hydrolase) | linear amide C-N hydrolisation | Bile salt hydrolase like bile resistance | *L. fermentum* | 220572-221498 | + |
| ***PpaC*, *L1970_00970***, Manganese-dependent inorganic pyrophosphatase (EC 3.6.1.1) | pyrophosphatase activity | Acid and bile resistance | *L. fermentum* | 191732-192664 | + |
| **Adhesion ability** | | | | | |
| **NFACT family protein** WP_236098355.1 | Fibronectin/fibrinogen-binding protein | Adhesion ability | *L. fermentum* | 517538-519223 | - |
| ***srtA*** (Sortase A, mucus specific LPXTG surface adhesin) | Surface proteins modification | Adhesion/Bile resistance | *L. fermentum* | 1422835-1423536 | + |
| **Psychobiotic function-related genes** | | | | | + |
| ***gadB, gadA*** **(**Glutamate decarboxylase (EC 4.1.1.15)) L1970_07645 | pyridoxal-dependent decarboxylase | γ-aminobutyric acid (GABA) synthesis | *L. fermentum* | 1555527-1556480 | - |
| ***gadC*** (glutamate: GABA antiporter) | GABA transporter | GABA release | *L. fermentum* | 1080245-1081537 | - |

**Table S16.** The bactericidal activity test results of the AGA52 against the test pathogens

| **Test Strain** | **Strain Code** | **Zone of Inbition (±SD)** |
|---|---|---|
| *Yersinia enterocolitica* | ATCC 9610 | 16,99 ± 0,44 |
| *Bacillus cereus* | ATCC 33019 | 7,47 ± 0,06 |
| *Salmonella enterica sv. Typhimurium* | ATCC 14028 | 7,6 ± 0,01 |
| *Escherichia coli O157:h7* | ATCC 43895 | 11,55 ± 0,09 |
| *Listeria monocytogenes* | ATCC 7644 | 11,33 ± 0,38 |
| *Klebsiella prenumoniae* | ATCC 13883 | 7,10 ± 0,2 |
| *Proteus vulgaris* | ATCC 8427 | 8,10 ± 0,17 |

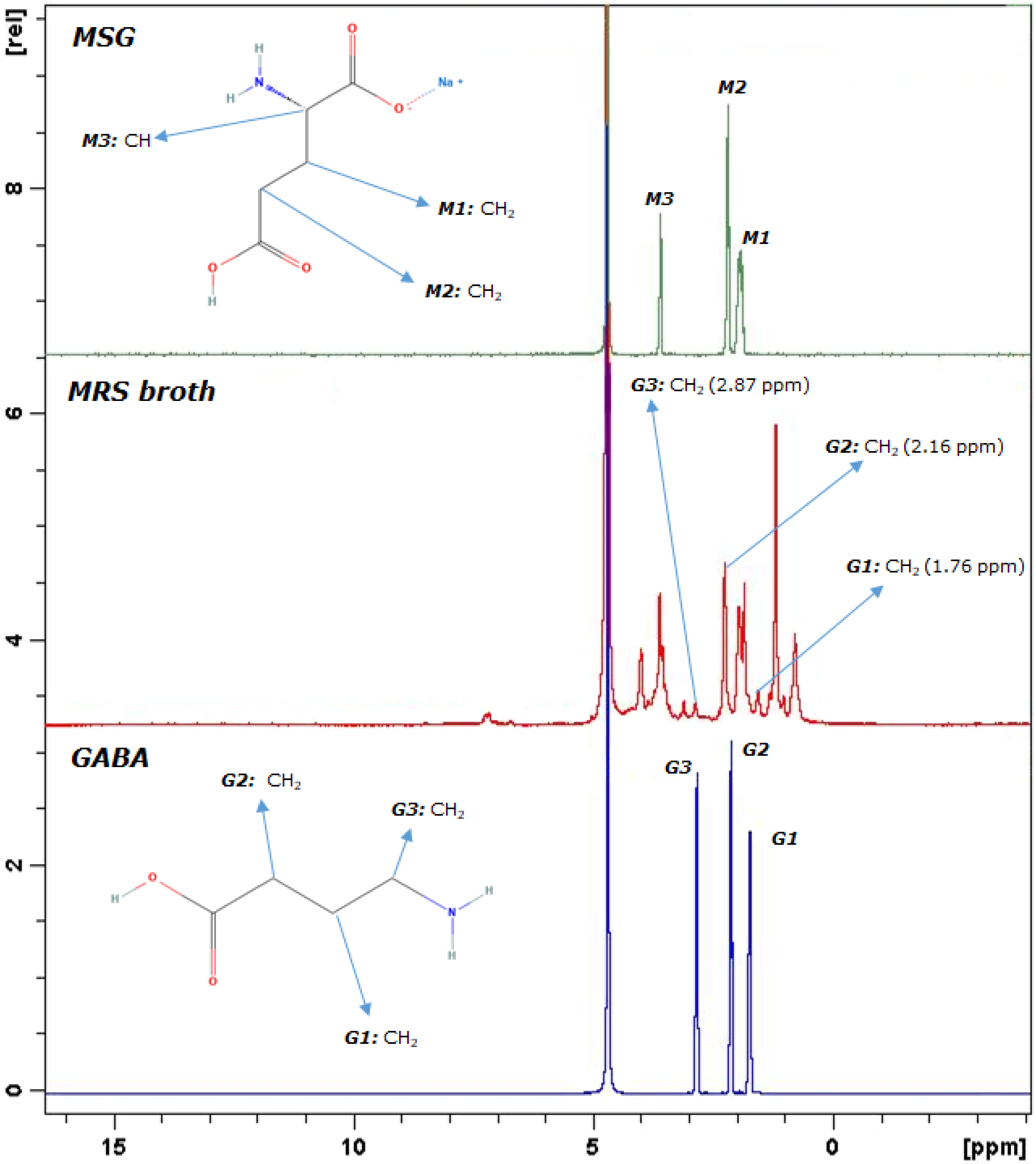


**Fig S4.** Confirmation of the GABA production by *Lb. fermentum* AGA52 in MRS broth environment by using NMR Spectra. Besides, standard measurements from monosodium glutamate (MSG) and γ-aminobutyric acid (GABA) were pefromed and both of carboxyl (-COOH) and amino ($-NH_2$) groups can't be found in their respective spectra due to zwitter-ion effect. M1: $CH_2$ (1.95 ppm), M2: $CH_2$ (2.2 ppm), M3: CH (3.6 ppm), G1: $CH_2$ (1.75 ppm), G2: $CH_2$ (2.15 ppm), and G3: $CH_2$ (2.85 ppm).
